# Effects of Environmental Salinity on Global and Endocrine-Specific Transcriptomic Profiles in the Caudal Neurosecretory System of Salmonid Fishes

**DOI:** 10.1101/2024.12.13.627795

**Authors:** Brett M. Culbert, Stephen D. McCormick, Nicholas J. Bernier

**Affiliations:** Department of Integrative Biology, University of Guelph, 50 Stone Rd E, Guelph, Ontario, Canada; Department of Biology, University of Massachusetts, Amherst, Amherst, MA, United States

**Keywords:** corticotropin-releasing hormone, neuropeptides, neurosecretory systems, *Oncorhynchus mykiss*, osmoregulation, *Salmo salar*, transcriptome

## Abstract

The caudal neurosecretory system (CNSS) is a fish-specific neuroendocrine complex whose function(s) remain uncertain despite 60+ years of research. Osmoregulatory roles for the CNSS have been hypothesized, but molecular regulation of the CNSS following changes in environmental salinity remains poorly characterized. Therefore, we performed transcriptomics on the CNSS of rainbow trout (*Oncorhynchus mykiss*) to establish: 1) how the CNSS responds following seawater (SW) transfer, and 2) which endocrine systems contribute to osmoregulatory responses in the CNSS. Responses following SW transfer varied at 24 h versus 168 h, with changes primarily affecting membrane transport and transcriptional processes at 24 h and neuronal processes at 168 h. Components of several osmoregulation-associated endocrine systems were affected (e.g., corticosteroid receptors), including some which have not previously been identified in the CNSS (e.g., calcitonin). Additionally, transcript levels of corticotropin-releasing factor (CRF) peptides—which have osmoregulatory functions and were highly abundant in the CNSS—were ∼2-fold higher after 24 h in SW. Therefore, we performed additional experiments investigating CRF peptides in a more euryhaline salmonid, Atlantic salmon (*Salmo salar*). Smolts had up to 12-fold higher levels of CRF peptide transcripts than parr, but abundance declined following SW transfer. Additionally, CRF transcripts were lower 24 h following freshwater transfer of SW-acclimated salmon. These results suggest that CRF peptides acutely aid in coordinating physiological responses following fluctuations in environmental salinity via anticipatory and/or responsive mechanisms. Collectively, our data indicate that CNSS-mediated production of CRF peptides has osmoregulatory functions and provide a resource for investigations of novel CNSS functions.

## 1. Introduction

The caudal neurosecretory system (CNSS) is a neuroendocrine complex that is located at the distal tip of the spinal cord in fishes. In teleosts, the CNSS is comprised of neurosecretory cells (i.e., Dahlgren cells) whose axons project towards capillary beds located in the caudal vertebrae and converge to form a neurohemal organ called the urophysis (Bern and Takasugi, 1962; Chan and Bern, 1976; Rousseau et al., 2024). The urophysis was first anatomically described in the early 1800s (Weber, 1827), but the presence of enlarged neurosecretory cells in the caudal spinal cord was not noted until about 100 years later (Dahlgren, 1914). Ultimately, these observations provided the groundwork for the later development of the CNSS as an endocrine complex (Enami, 1959). While the importance of the CNSS in the field of endocrinology is cemented owing to the eponymous naming of two peptide hormones that were first identified in and isolated from the CNSS—a peptide in the corticotropin-releasing factor (CRF) family [urotensin 1, UTS1; (Lederis et al., 1982)] and a peptide in the somatostatin family [urotensin 2, UTS2; (Pearson et al., 1980)]—specific functional roles for this unique neuroendocrine complex remain unclear 60+ years later.

The CNSS has been hypothesized to contribute to several physiological processes, including reproduction, stress responses, cardiovascular regulation, and thermal adaptation (Chan and Bern, 1976; Munro, 1995; Yuan et al., 2020a). However, the greatest attention has been given to the role of the CNSS during changes in environmental salinity. Early studies noted that the size and content of neurosecretory cells in the CNSS changed following changes in salinity (Chevalier, 1976; Enami, 1956; Takasugi and Bern, 1962), and removal of the CNSS reduced the ability of some species to acclimate to altered salinities (Ireland, 1969; Takasugi and Bern, 1962). These osmoregulatory actions were hypothesized to be mediated by products synthesized within the CNSS because injection of urophysial extracts induced osmoregulatory responses (Enami et al., 1956; Fryer et al., 1978; Takasugi and Bern, 1962). Furthermore, changes in either the external (Ashworth et al., 2005; Yagi and Bern, 1963) or internal (Bennett and Fox, 1962; Yagi and Bern, 1965) osmotic environment affect the electrical activity of neurons within the CNSS. However, responses vary depending on changes in specific ions. For example, neurons in the CNSS of tilapia (*Oreochromis mossambicus*) can be separated into those that increase firing rates when exposed to high Na^+^ and those that respond when Na^+^ levels are low, but responses to Na^+^ levels appear to be regulated by the brain and not the CNSS directly (Yagi and Bern, 1965). In contrast, the presence of Ca^+2^ sensing receptors on Dahlgren cells (Greenwood et al., 2009; Ingleton et al., 2002), as well as high transcript levels of the calcium-regulating hormones parathyroid hormone-related peptide [PTHrP; (Ingleton et al., 2002; Lu et al., 2017)] and stanniocalcin-1 [STC-1; (Greenwood et al., 2009)] in the CNSS of flounder (*Platichthys flesus*) suggests that the CNSS can directly sense changes in internal Ca^+2^ levels and participates in Ca^+2^ regulation. Yet, the generalizability of these findings is limited because most work on the CNSS has focused on only a handful of species. Furthermore, despite a solid understanding of the electrophysiological responses displayed by the CNSS in response to changes in environmental salinity, transcriptional regulation of the CNSS following salinity changes is still poorly understood.

Many of the osmoregulatory effects described above were later attributed to changes in production of UTS1 and/or UTS2 by the CNSS (Arnold-Reed et al., 1991; Minniti et al., 1989). However, changes in the synthesis and release of these hormones by the CNSS—as well as specific osmoregulatory actions of UTS1 and UTS2 across different epithelial tissues (Chan, 1975; Loretz et al., 1981; Mainoya and Bern, 1982; Marshall and Bern, 1981)—often vary among species, likely reflecting species-specific osmoregulatory ability and interactions with other hormone and neurotransmitter systems. Indeed, in addition to synthesizing UTS1 and UTS2, the CNSS also contains large amounts of CRFb (Alderman and Bernier, 2009; Craig et al., 2005; Lu et al., 2004) and PTHrP (Ingleton et al., 2002; Lu et al., 2017), as well as smaller amounts of arginine vasotocin/vasopressin (AVP), isotocin/oxytocin, and urocortin’s 2 & 3 in some species (Gozdowska et al., 2013; Grone et al., 2021; Harding et al., 1997; Holder et al., 1979; Hosono et al., 2017; Lacanilao, 1972; Parmentier et al., 2008). Furthermore, the presence of descending peptidergic nerve fibers—as well as cholinergic, adrenergic, and serotonergic fibers (Audet and Chevalier, 1981; Conlon and Balment, 1996; Hubbard et al., 1996; McKeon et al., 1988; Yulis et al., 1990)—that exhibit positive staining for GnRH (Miller and Kriebel, 1986; Xia et al., 2015; Yamamoto et al., 1995) and NPY (Oka et al., 1997), as well as receptors for CRF and UTS1, UTS2, cortisol, prolactin, and GnRH (Lu et al., 2007) indicate that the CNSS is likely a major endocrine player in fishes. However, the full complement of endocrine factors produced in the CNSS—and the suite of hormone systems which contribute to the regulation of the CNSS—remains uncertain.

In the current study, we conducted the first transcriptomic assessment of how the CNSS responds following changes in environmental salinity. Specifically, we evaluated changes either 24 or 168 h following transfer of rainbow trout (*Oncorhynchus mykiss*) from freshwater (FW) to seawater (SW). We used rainbow trout because they are euryhaline, and previous work has shown that the CNSS is responsive to osmoregulatory changes in rainbow trout and other salmonids (Chevalier, 1976; Craig et al., 2005; Larson and Madani, 1996; Nishioka et al., 1982). Another major focus of the current study was to identify which components of different endocrine systems are present in the transcriptome of the rainbow trout CNSS to shed light on potential novel endocrine functions and/or regulators of this tissue, especially those involved in osmoregulatory processes. While a previous study (Yuan et al., 2021) used a transcriptomics approach to evaluate whether responses of the CNSS following a temperature change varied between bold and shy olive flounder (*Paralichthys olivaceus*), they did not describe the transcriptome of the CNSS in detail. Thus, the full complement of endocrine systems present in the CNSS remains unclear. We also performed several additional experiments to determine how environmental salinity affects transcript levels of CRF peptides (specifically, CRFb and UTS1) in the CNSS. We focused on the CRF system because: 1) CRF peptides are abundant in the CNSS (Bernier et al., 2008; Lu et al., 2004); 2) transcript and protein levels of CRF peptides in the CNSS often vary following changes in environmental salinity (Craig et al., 2005; Lu et al., 2004; Zhu et al., 2019); and 3) the CRF system has conserved osmoregulatory functions (Cannell et al., 2016; Loretz et al., 1981; Mainoya and Bern, 1982; Marshall and Bern, 1981; Stengel and Taché, 2009). For some of these experiments, we used Atlantic salmon (*Salmo salar*) because they are anadromous [i.e., migrate between FW and SW as part of their natural life history (McCormick, 2012; Zydlewski and Wilkie, 2012)], offering the opportunity to evaluate how CRF peptide synthesis in the CNSS is affected by environmental salinity under more ecologically relevant conditions. Specifically, we investigated how transcript levels of CRF peptides changed: 1) during the seasonal acquisition of SW tolerance as salmon transform from pre-migratory parr into migratory smolts (i.e., smoltification); 2) in response to SW transfer of FW-acclimated parr and smolts; and 3) in response to FW transfer of SW-acclimated post-smolts.

## 2. Materials and Methods

### 2.1. Experimental Animals and Housing

Experiments 1 and 3 took place at the Hagen Aqualab at the University of Guelph (Guelph, ON, Canada) using either rainbow trout (Exp. 1) or Atlantic salmon (Exp 3) that were ∼3 years old. Rainbow trout were acquired from the Ontario Aquaculture Research Centre (Alma, ON, Canada), while Atlantic salmon were acquired from the Normandale Fish Culture Station (Vittoria, ON, Canada). Both species were maintained in 1.8 m diameter fibreglass tanks (∼2000L) that were supplied with aerated, flow-through well water at 12°C and kept on a 12h light, 12h dark photoperiod regime. Fish were fed to satiation 3 times per week with commercial pellets (Blue Water Fish Food; Guelph, ON, Canada). A stocking density of ∼100 fish per tank was maintained and fish were kept under these conditions for several months prior to starting experiments. All procedures were carried out in accordance with the Canadian Council for Animal Care guidelines for the use of animals in research and teaching and were approved by the University of Guelph’s Animal Care Committee (AUP #4123).

Experiment 2 took place at the U.S. Geological Survey (USGS) S.O. Conte Anadromous Fish Research Laboratory (Turners Falls, MA, USA) using ∼1 year old juvenile Atlantic salmon that were obtained from the Kensington State Hatchery (Kensington, CT, USA) in the Fall of 2018. Fish were held in 1.8 m diameter tanks that were supplied with flow-through ambient Connecticut River water at a flow rate of 4 L min^−1^, and tanks were continuously aerated. Fish were maintained under the natural photoperiod and were fed to satiation (BioOregon; Westbrook, ME, USA) using automatic feeders. In December of 2018, fish were separated by size into parr and pre-smolt groups as described previously (Culbert et al., 2022). Each group of fish was maintained in duplicate tanks containing ∼100 fish, and all fish experienced identical temperature regimes throughout the experiment. All fish rearing and sampling protocols were carried out in accordance with USGS institutional guidelines and protocol LSC-9096, which was approved by the USGS Eastern Ecological Science Center Institutional Animal Care and Use Committee.

### 2.2. Experiment 1: Regulation of the CNSS following FW-to-SW transfer of rainbow trout

#### 2.2.1. Experimental Details

Fifty-eight trout (mass = 545 ± 14 g; fork length = 35.0 ± 0.3 cm; mean ± SEM) were moved from the stock tank and randomly placed into one of six 2000L recirculating tanks (N=8-10 fish per tank). Three tanks contained well water (FW) and three contained SW (35ppt, Instant Ocean Sea Salt; Blacksburg, VA, USA). Each tank received continuous aeration via an air stone and was equipped with both particle and biological filtration, as well as UV sterilization. Groups of fish were sampled either 24, 72, or 168 h following transfer and no mortalities occurred. Food was withheld throughout the experiment because salmonids naturally suppress food intake following transfer to SW (Arnesen et al., 1993; Craig et al., 2005; Damsgård and Arnesen, 1998; Usher et al., 1991). All fish were killed between 1000 and 1200 h via terminal anaesthesia using buffered MS-222 (100 mg L^-1^, Syndel; Vancouver, BC, Canada). Fork length and mass were recorded, blood was collected from the caudal vasculature using a 1 mL ammonium heparinized syringe and was spun at 9,000 *g* for 5 min. Most fish (>80%) were still juvenile and therefore sex was not included in our analyses. After this, plasma was collected, frozen on dry ice, and stored at -80°C for later determination of osmolality and cortisol levels. To sample the CNSS, we collected all spinal tissue from the five most caudal vertebrae and the urostyle. Following dissection, the CNSS was frozen on dry ice, and stored at -80°C for later extraction of RNA (see section 2.5) to be used for RNA-Seq (see section 2.7) and quantitative polymerase chain reaction (qPCR; see section 2.6). A portion of the CNSS was lost while dissecting one of the 7 d SW-transferred fish, resulting in N = 7 for this group.

### 2.3. Experiment 2: Transcriptional regulation of CRF peptides in the CNSS during smoltification and FW‒to-SW transfer of juvenile Atlantic salmon

#### 2.3.1. Experimental Details

Twelve parr and twelve smolts were randomly sampled on February 19^th^, April 1^st^, and May 6^th^ in 2019. Fish were kept at ambient temperatures (2-4°C) through the winter and water temperature was increased by 1°C per day until 8-10°C was reached on February 15. Temperature was kept the same throughout the experiment so that all fish were sampled at identical temperatures. To avoid potential tank effects, fish from each group (parr or smolts) were sampled from two separate tanks at each timepoint (N=6 per tank). To determine the response of fish following SW exposure, groups of parr and smolts were randomly selected and placed into six 1 m diameter tanks (3 tanks of parr and 3 tanks of smolts) containing 28 ppt recirculating SW (Instant Ocean Sea Salt) during the week of May 6th, 2019, which is the peak of smolt development for this population under these conditions (McCormick et al., 2013). These tanks were held at 8.5-9.5°C and contained particle, biological, and charcoal filtration, as well as continuous aeration. Fish were fed to satiation every day, but food was withheld the day prior to samplings. Twelve parr and twelve smolts were sampled after 24, 96, and 240 h of SW exposure and no mortalities occurred. All fish were killed via terminal anesthesia using buffered MS-222 (100 mg L^−1^) after which fork length and mass were recorded. All fish were juvenile and therefore sex could not be determined. Blood was collected from the caudal vasculature using a 1 mL ammonium heparinized syringe, spun at 3,200 g for 5 min at 4°C, and plasma was collected for later measurement of cortisol and osmolality. Given the small size of these fish, the CNSS was fixed prior to removal from the spinal column to prevent tissue loss. All muscle was removed from the caudal spinal column (consisting of the five most caudal vertebrae and the urostyle) and the spinal column was placed in a solution of 4% paraformaldehyde (PFA) in phosphate-buffered saline (PBS; pH 7.4) for 24 h at 4°C. The following day, the CNSS was removed from the spinal column and RNA was extracted following the protocol of Craig et al. (2005) for later qPCR analysis (see section 2.6).

### 2.4. Experiment 3: Transcriptional regulation of CRF peptides in the CNSS during SW‒to-FW transfer of post-smolt Atlantic salmon

#### 2.4.1. Experimental Details

Salmon (N= 28; fork length: 37.8 ± 0.6 cm, mass: 537.8 ± 24.0 g) were held in a recirculating tank which contained ∼2000L of SW (33ppt; Instant Ocean Sea Salt) for approximately 6 months. This tank was continuously aerated with an air stone and was equipped with both particle and biological filtration, as well as UV sterilization. At the start of the experiment, N=10 salmon were immediately sampled while the remaining fish were randomly split among two ∼500L tanks containing aerated, flow-through well water (FW; N=9 per tank). After either 24 h or 96 h in FW, fish were terminally anesthetized using phenoxyethanol (0.2%) and their plasma (for cortisol and osmolality) and CNSS (for qPCR; see section 2.6) were collected as previously described. We also sampled FW-acclimated salmon (N=10; fork length: 39.4 ± 1.0 cm, mass: 581.3 ± 42.3 g) for comparison (see section 2.1 for housing conditions). No mortalities occurred following FW transfer and all treatment groups contained approximately equal proportions of females and males (53% male and 47% female combined).

### 2.5. Plasma Cortisol and Osmolality Measurements

Plasma osmolality values were determined in duplicate using a Vapro 5520 vapour pressure osmometer (Wescor; Logan, UT, USA) and had an intra-assay variation of 8.3% coefficient of variation (CV). Circulating cortisol levels were determined using a commercially available enzyme immunoassay (Neogen, Cat # 402710; Lexington, KY, USA) for experiments 1 and 3, or a previously validated direct competitive enzyme immunoassay (Carey and McCormick, 1998) for experiment 2. Intra- and inter-assay variation was 8.2% and 13.1% CV, respectively.

### 2.6. RNA Isolation and qPCR

The entire CNSS of fish from experiments 1 and 3 was homogenized in TRIzol reagent (Invitrogen; Burlington, ON, Canada) using a Precellys Evolution tissue homogenizer (Bertin Instruments; Montigny-le-Bretonneux, France) and total RNA was extracted as per the manufacturer’s directions (see Craig et al., 2005 for details of RNA extraction of CNSS samples from experiment 2). The quantity and purity of RNA samples was assessed using a NanoDrop 2000 spectrophotometer (Thermo Scientific; Mississauga, ON, Canada). Following this, we treated 1µg of RNA with DNase (DNase 1; Thermo Scientific) and reverse transcribed cDNA using a high-capacity Applied Biosystems cDNA reverse transcription kit (Thermo Scientific). We then performed qPCR using a CFX96 system (BioRad; Hercules, CA, USA) with SYBR green (SsoAdvanced Universal; BioRad) and gene-specific primers (Table 1). All samples were run in duplicate and negative controls—including no template controls (where cDNA was replaced with water) and no reverse transcriptase controls (where RNA reverse transcriptase was replaced with water during cDNA synthesis)—were also included. Each reaction contained a total of 20 µl, which consisted of 10 µl of SYBR green, 5 µl of combined forward and reverse primers (0.2 µM [final]), and 5 µl of 10x diluted cDNA. Cycling parameters included a 30 s activation step at 95°C, followed by 40 cycles consisting of a 3 s denaturation step at 95°C and a combined 30 s annealing and extension step at 60°C. Melt curve analysis was conducted at the end of each run to confirm the specificity of each reaction. To account for differences in amplification efficiency, standard curves were constructed for each gene using serial dilutions (4x) of pooled cDNA. Input values for each gene were obtained by fitting the average threshold cycle value to the antilog of the gene-specific standard curve, thereby correcting for differences in primer amplification efficiency. To correct for minor variations in template input and transcriptional efficiency, we normalized our data to the geometric mean of transcript abundances of elongation factor 1α (*ef1α*) and ribosomal protein L13a (*rpl13a*) as reference genes. While reference genes were stable across treatments in experiments 1 and 3, expression of reference genes varied between sampling times in experiment 2. To account for differences in reference gene expression across time, the geometric mean of the reference genes within each group of samples was normalized to a “control” group (parr in February) using a correction factor (mean value within a group/mean value of control group) as previously described (Billiau et al., 2001; Essex-Fraser et al., 2005). All data are expressed relative to the mean value of the control group within each experiment (see figure captions for further details).

**Table 1.**
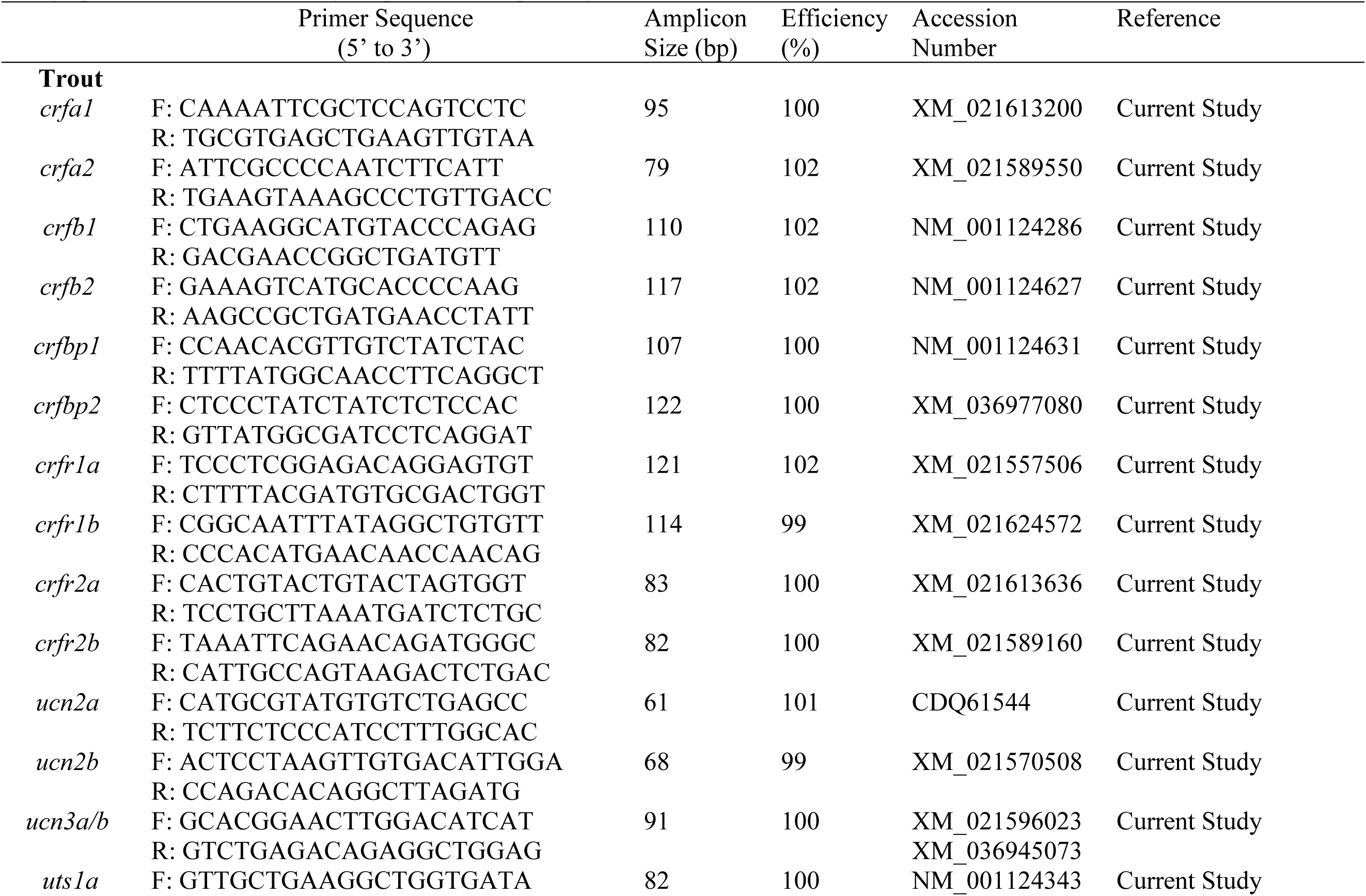
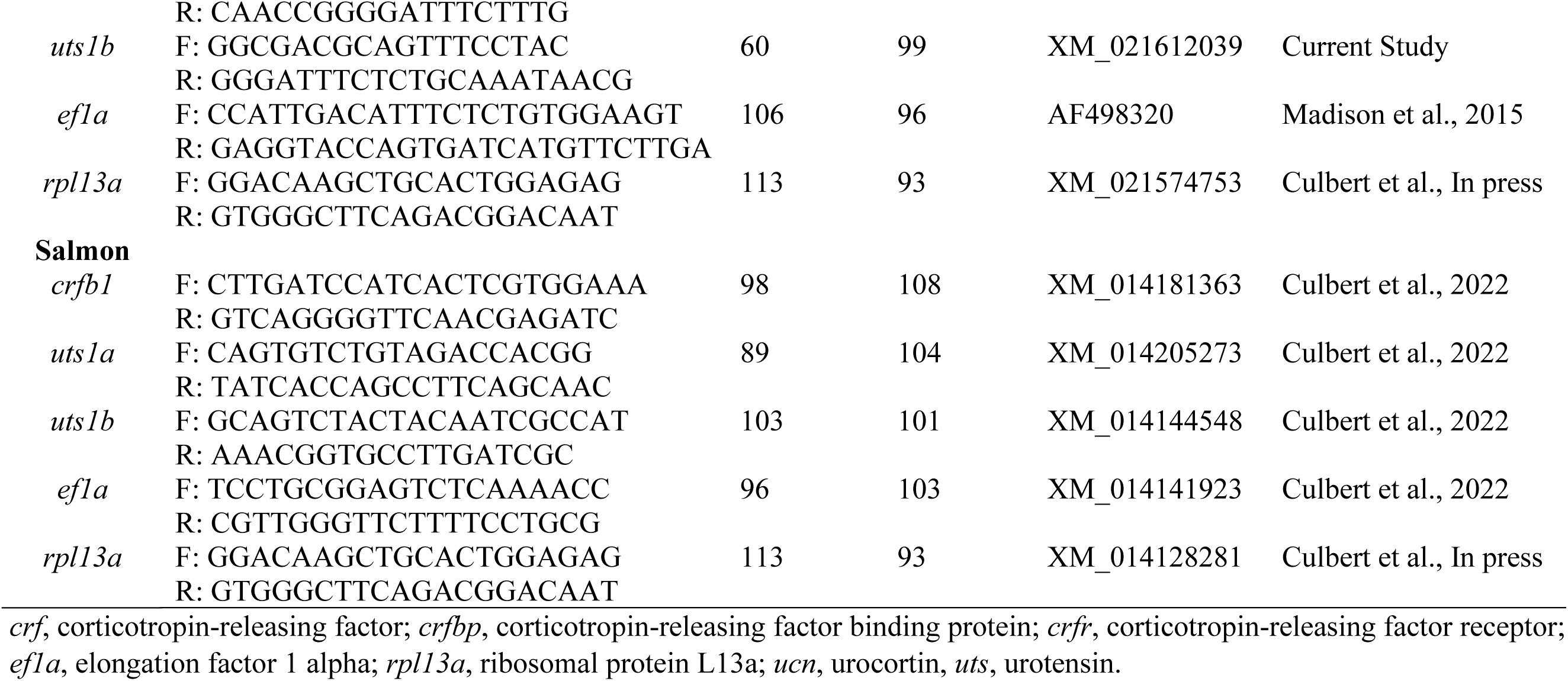
Gene specific primers used for real-time polymerase chain reaction (qPCR) in rainbow trout (*Oncorhynchus mykiss*) or Atlantic Salmon (*Salmo salar*). Note that the sequences of *ucn3a* and *ucn3b* are 99% identical in both species and we were therefore unable to design primers that differentially amplified these paralogs.

### 2.7. RNA-Seq

To evaluate both acute and chronic responses of the rainbow trout CNSS during SW acclimation, we conducted RNA-Seq on FW-to-FW (control) and FW-to-SW transferred fish (N=7-9 per group) either 24 or 168 h post-transfer. Total RNA (see section 2.6) was sent to the Génome Québec Centre of Expertise and Services (Montreal, QC, Canada) for library preparation and sequencing. The integrity of all RNA was evaluated using a Bioanalyzer 2100 (Agilent; Mississauga, ON, Canada) and all samples had RNA Integrity Numbers (RIN) greater than 7.6 (8.1 ± 0.3). Using 250ng of total RNA, mRNA was isolated using NEBNext® Poly(A) Magnetic Isolation Module (New England Biolabs; Whitby, ON, Canada). Stranded cDNA libraries [296 ± 1 base pair (bp) length] were created using the Illumina Stranded mRNA Prep and sequencing was conducted on all 32 samples using 100 bp paired-end reads with a NovaSeq 6000 (Illumina; San Diego, CA, USA). Samples were sequenced across two batches with equal representation from each group within each run. A mean of ∼35,000,000 ± 4,000,000 paired-end reads were produced per sample (Supp. Table 1). Preliminary analyses indicated that batch effects were negligible (<1% of variation) and therefore batch was not included in statistical models. Raw sequencing reads are archived with the National Center for Biotechnology Information Sequence Read Archive (NCBI SRA Accession: PRJNA1139945).

Processing and analysis of the RNA-Seq data was performed by the Canadian Centre for Computational Genomics (C3G; https://computationalgenomics.ca/) using their in-house RNA-Seq pipeline (https://bitbucket.org/mugqic/genpipes/src/master/pipelines/rnaseq/). Briefly, paired reads were joined and checked for quality using FastQC (v0.12.0, RRID:SCR_014583; Babraham Bioinformatics) and low-quality reads (Phred score < 30; ∼0.02% of all reads) were trimmed using Trimmomatic [v0.39.0, RRID:SCR_011848; (Bolger et al., 2014)]. Filtered reads from all libraries were aligned to the rainbow trout reference genome (USDA_OmykA_1.1; ENSEMBL version 107; NCBI Accession: GCA_013265735.3) using STAR [v2.7.8a, RRID:SCR_004463; (Dobin et al., 2013)]. This resulted in an average alignment rate of 86.3 ± 1.2 % across all samples which provided an average of 12,934 ± 78 transcripts per sample. A total of 49,009 unique transcripts were identified overall. Picard (v3.0.0, RRID:SCR_006525; Picard Tools) was used to create a single global BAM file for each sample and RNA expression quantification was performed using HTSeq-count [v2.0.2, RRID:SCR_011867; (Anders et al., 2015)]. Using the edgeR package [v3.40.2, RRID:SCR_012802; (Robinson et al., 2009)] in R [v4.2.3, RRID:SCR_001905; (R Core Team, 2023)], genes were filtered for low expression (a counts per million score of ≥ 3) which resulted in a final count of 25,647 unique transcripts.

### 2.8. Statistical Analyses

For all RNA-Seq analyses, each sample was normalized to its overall library size and differential gene expression testing was performed in edgeR [v3.42.4, RRID:SCR_012802; (Robinson et al., 2009)] using generalized linear models and F-tests. For the main RNA-Seq analysis, the resulting p-values from these analyses were false-discovery rate corrected (FDR; p < 0.01) using the Benjamini-Hochberg (BH) method within edgeR and an effect size of log_2_(fold change) >|0.5| between groups was utilized. However, we used a more liberal FDR threshold of p < 0.05 without a fold change threshold when exploring changes in endocrine genes. Functional enrichment analysis and subsequent data visualization was performed as described by Reimand et al. (2019). Briefly, Gene Ontology (GO) terms were generated from gene lists via the g:GOSt function within g:Profiler [RRID:SCR_006809; (Raudvere et al., 2019)]. GO terms were considered as significantly enriched if they contained 4 or more differentially expressed genes (DEGs) and had a BH-FDR value ≤ 0.1. To account for the identification of multiple GO terms that all contained the same sets of DEGs, we created GO clusters using the EnrichmentMap [v3.3.6, RRID:SCR_016052; (Merico et al., 2010)] and AutoAnnotate [v1.4.0;(Kucera et al., 2016)] apps in Cytoscape [v3.10.0, RRID:SCR_003032; (Shannon et al., 2003)]. GO clusters were annotated according to words that were most common across all GO terms within each cluster. Enrichment maps were created using each list of significant GO terms and their respective clusters. Additionally, to better characterize the extent of overlap between DEGs across groups, we also created upset plots using the ‘UpSetR’ package (Conway et al., 2017) and biplots depicting principal components 1 and 2—which were derived from a principal components analysis using the ‘prcomp’ function—using the ‘ggbiplot’ package.

Statistical analysis of all plasma and qPCR data was performed using R [version 4.3; R Core Team, 2023)]. All data are presented as means ± 1 standard error of the mean (SEM) and a significance level (α) of 0.05 was used for all tests. Outliers were excluded based on a 2x interquartile range threshold. When data did not meet the assumptions of normality and/or equal variance, data were either log or square-root transformed to meet the assumptions, or analyses were performed using ranked data. Plasma and qPCR data for Experiments 1 and 2 were analyzed using two-way ANOVAs that included group (FW and SW or parr and smolt) and either month (February, April or May) or time following SW exposure (FW, 24 h, 96 h or 240 h), as well as the interaction between these factors. Data from Experiment 3 were analyzed using one-way ANOVAs with time as a fixed factor. When significant differences were detected, post hoc Tukey’s tests were performed using the ‘emmeans’ package (Lenth, 2016).

## 3. Results

### 3.1 Experiment Series 1: Rainbow Trout

#### 3.1.1 Physiological effects of FW-to-SW transfer

Plasma osmolality levels (Fig. 1A; p_group_<0.001, p_time_=0.07, p_group*time_=0.11) were ∼25% higher in FW-to-SW transferred fish compared to FW-to-FW transferred fish across all timepoints. In contrast, while plasma cortisol levels (Fig. 1B; p_group_<0.001, p_time_=0.16, p_group*time_<0.001) were low across all timepoints in FW-to-FW transferred fish, cortisol levels in FW-to-SW transferred fish were 3- and 16-fold higher 24 and 72 h post-transfer, respectively.

**Figure 1.**
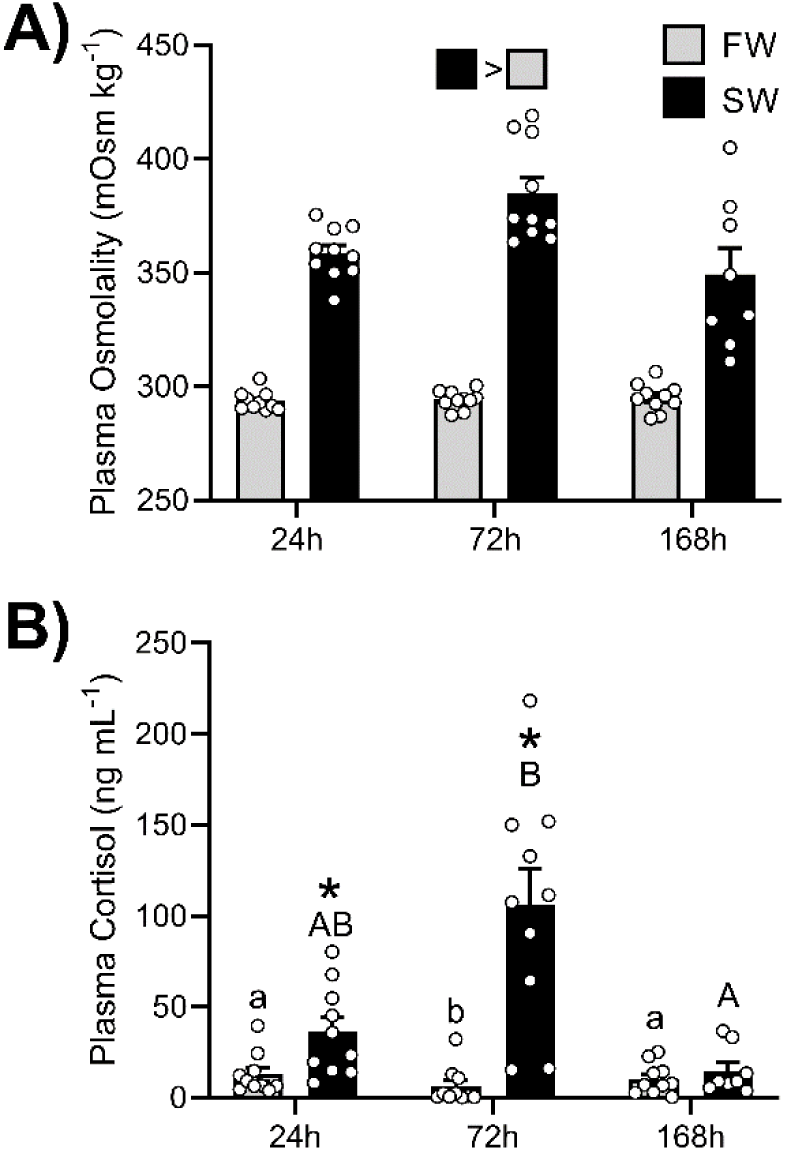
Changes in (A) plasma osmolality and (B) cortisol values of rainbow trout (*Oncorhynchus mykiss*) that were transferred from freshwater-to-freshwater (FW; grey) or from freshwater-to-seawater (SW; black) for 24, 72, or 168h. Significant differences (p < 0.05) are depicted using either letters (across time; uppercase = within SW, lowercase = within FW), filled oversized squares (between groups across all timepoints) or asterisks (between groups within a timepoint). Values are represented as means ± SEM and individual data points are shown.

#### 3.1.2 Effects of FW-to-SW transfer on CNSS transcriptome

After 24 h, we detected 324 DEGs (179 upregulated and 145 downregulated) when comparing FW-to-SW and FW-to-FW transferred fish (Supp. File 1). Further scrutiny of these DEGs using GO analysis revealed changes in clusters related to nuclear transport, membrane transport, DNA maintenance, negative transcription regulation, neurotransmitter transport, lipid biosynthesis, RAS protein signalling, and chromosome organization (Fig. 2; Supp. File 2). When only DEGs that were higher in the FW-to-SW group were included in the GO analysis neurotransmitter transport and stimuli responses differed (Fig. 2; Supp. File 2). In contrast, membrane transport, chromosome organization, and lipid biosynthesis were affected when GO analysis was performed using DEGs that were lower in the FW-to-SW group (Fig. 2; Supp. File 2).

**Figure 2.**
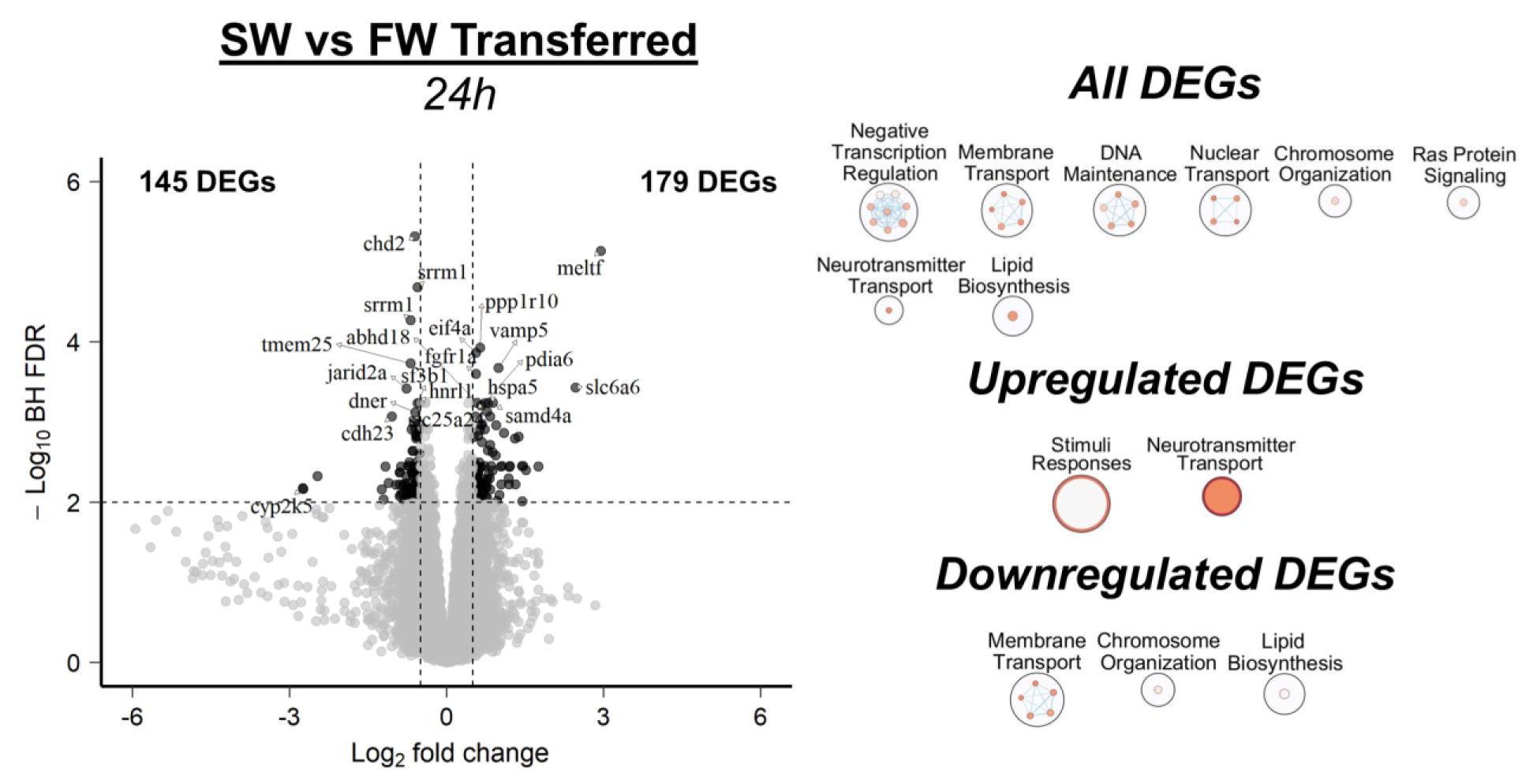
Effect of transferring rainbow trout (*Oncorhynchus mykiss*) from freshwater-to-freshwater (FW) or from freshwater-to-seawater (SW) for 24h as determined via RNA-Seq. Data are presented as changes in SW relative to FW. Differentially expressed genes (DEGs) with a Log_2_ fold change ≥ |0.5| between groups and were significantly different following Benjamini-Hochberg false-discovery rate (BH-FDR) correction (p < 0.01) are darkened on the volcano plot and a subset of genes are labelled (see Supplemental File 1 for more details). Note that for visualization purposes the x- and y- axes of the volcano plot are plotted on a Log_2_ and Log_10_ scale, respectively. Significant GO terms were clustered (larger circles) based on overlap between shared genes when analyzing all significant genes, only significant upregulated genes, or only significant downregulated genes. Each smaller circle within the larger circles represents an individual GO term (see Supplemental File 2 for more details). The extent of overlapping genes shared between GO terms is indicated by the intensity of the connecting lines (darker = more shared genes) and significance of each GO term is indicated by the colour of the circle (darker = more significant).

After fish had acclimated for 168 h, we detected 193 DEGs (96 upregulated and 97 downregulated) between FW-to-SW and FW-to-FW transferred fish (Supp. File 1). GO analysis revealed that these changes were primarily related to neuron development and differentiation, cell responses to chemical stimulus, protein folding, and phospholipid metabolism (Fig. 3; Supp. File 2). Specifically, upregulated DEGs were related to protein folding, while downregulated DEGs were related to neuron development and differentiation (Fig. 3; Supp. File 2).

**Figure 3.**
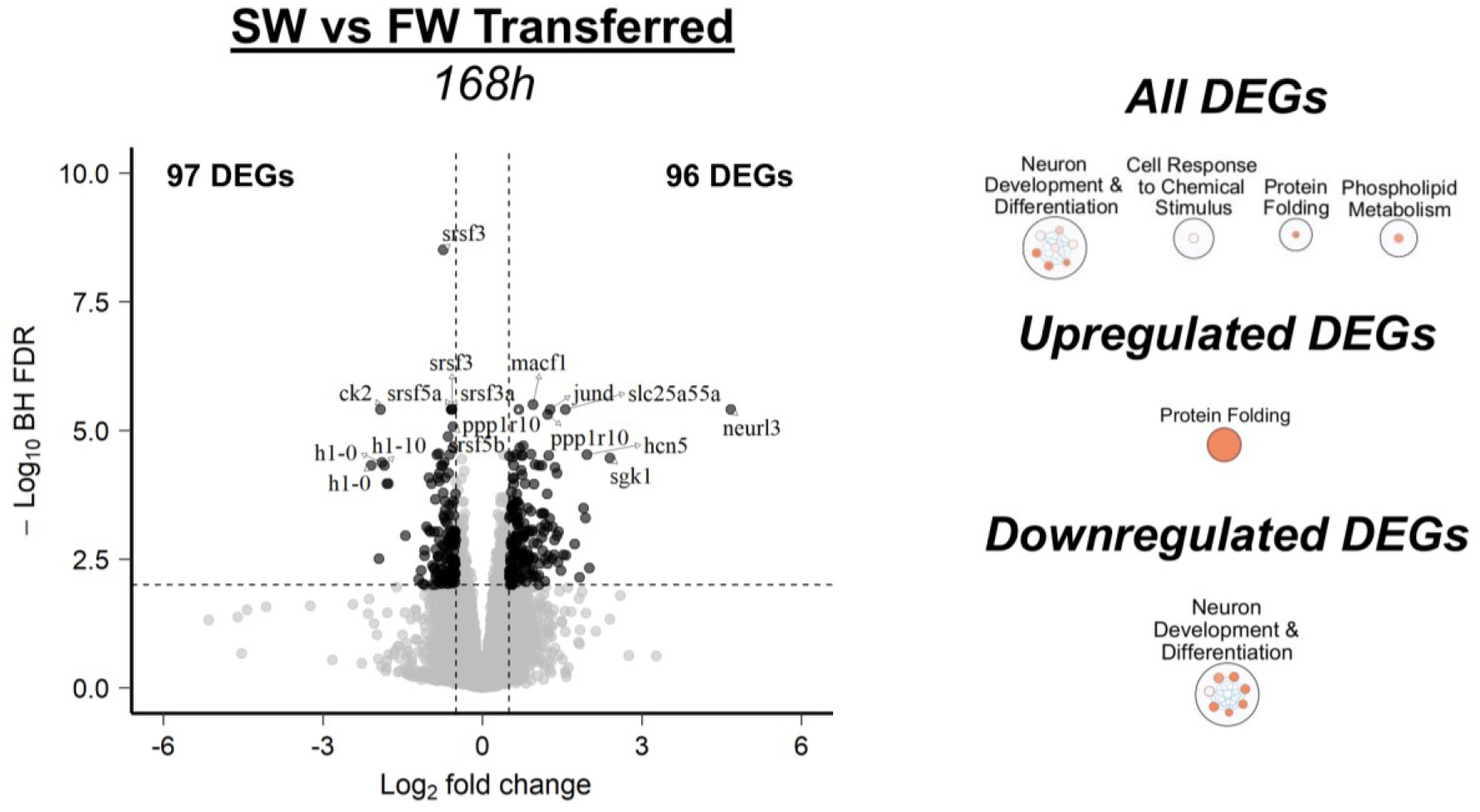
Effect of transferring rainbow trout (*Oncorhynchus mykiss*) from freshwater-to-freshwater (FW) or from freshwater-to-seawater (SW) for 168h as determined via RNA-Seq. Data are presented as changes in SW relative to FW. Differentially expressed genes (DEGs) with a Log_2_ fold change ≥ |0.5| between groups and were significantly different following Benjamini-Hochberg false-discovery rate (BH-FDR) correction (p < 0.01) are darkened on the volcano plot and a subset of genes are labelled (see Supplemental File 1 for more details). Note that for visualization purposes the x- and y- axes of the volcano plot are plotted on a Log_2_ and Log_10_ scale, respectively. Significant GO terms were clustered (larger circles) based on overlap between shared genes when analyzing all significant genes, only significant upregulated genes, or only significant downregulated genes. Each smaller circle within the larger circles represents an individual GO term (see Supplemental File 2 for more details). The extent of overlapping genes shared between GO terms is indicated by the intensity of the connecting lines (darker = more shared genes) and significance of each GO term is indicated by the colour of the circle (darker = more significant).

The greatest changes were observed when FW-to-SW transferred fish at 168 h versus 24 h were compared, which yielded 823 DEGs (425 upregulated and 398 downregulated; Supp. File 1). Accordingly, 17 unique DEG clusters were identified (Fig. 4; Supp. File 2), with several focusing on the regulation of DNA and RNA processing (cellular catabolism regulation, RNA catabolism, DNA organization and replication, RNA processing, and chromatin organization) and cellular signalling and synapse regulation (synaptic signalling, negative signalling and communication, and cell and synapse regulation). Additional GO clusters included: developmental processes, cell and membrane adhesion, cell proliferation regulation, negative cellular organization, amine metabolism, protein folding, cell projection organization, amino acid metabolism, and cell cycle regulation. When only upregulated DEGs were considered (Fig. 4; Supp. File 2), we again detected several GO clusters related to the regulation of DNA and RNA processing (DNA organization and replication, RNA processing, protein-DNA organization, DNA repair), as well as cell cycle regulation, negative biosynthesis regulation, cell projection organization, cellular catabolism regulation, glycoprotein regulation, tissue development, cell-cell adhesion, protein folding, amine metabolism, microtubule based movement, and amino acid metabolism. When focusing on downregulated DEGs (Fig. 4; Supp. File 2) several clusters related to synapses and cell proliferation (synapse signalling, cell proliferation, junction and synapse organization, and neuron generation), as well as actin filament regulation, RNA catabolism, negative cell signalling, substance and stimuli response, movement and responses, cell and membrane adhesion, cellular export and secretion, and protein dephosphorylation were identified. In contrast, we did not detect any significant DEGs when comparing the groups of fish that had been sampled 1- or 7-days following transfer from FW-to-FW (Supp. File 1).

**Figure 4.**
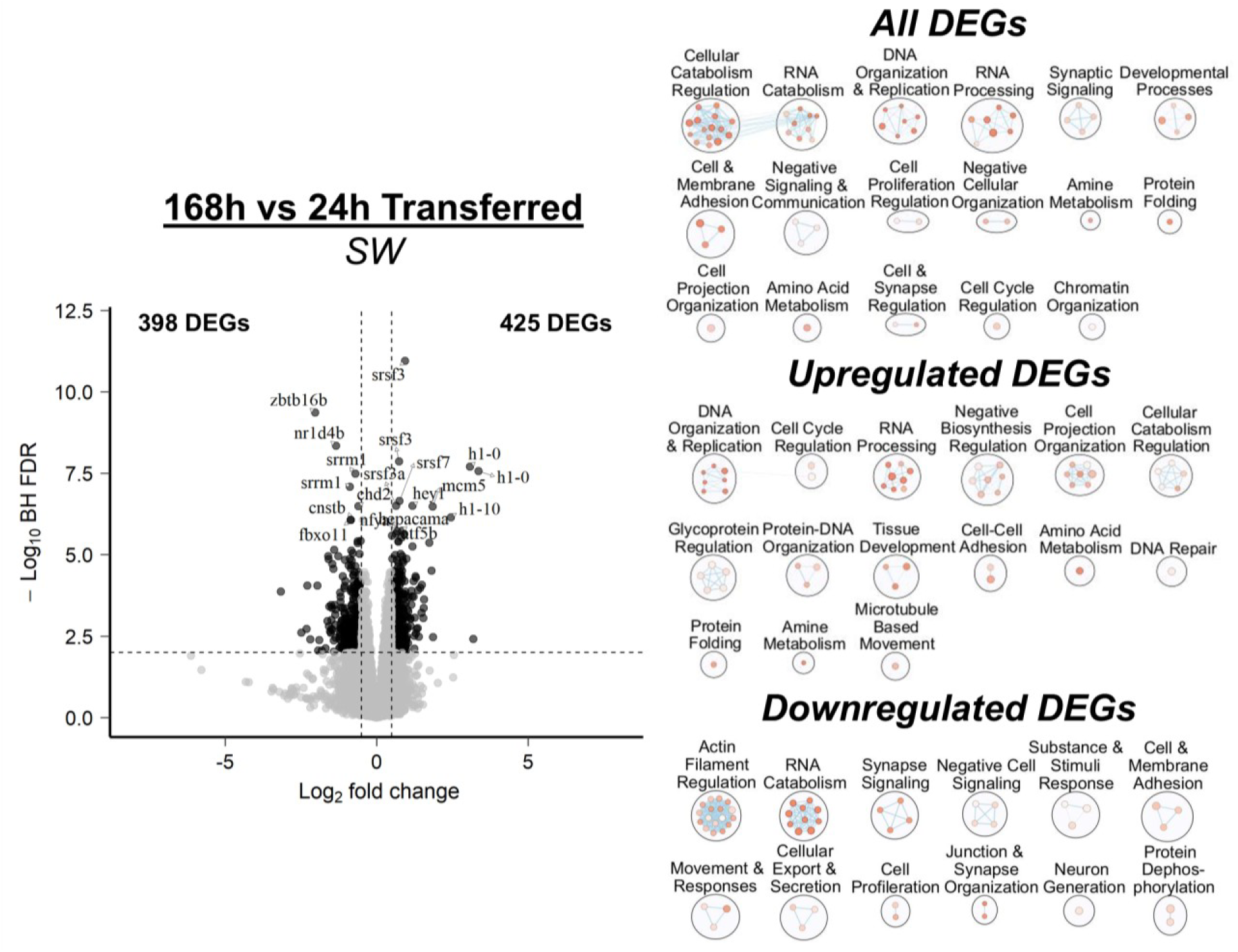
Effect of transferring rainbow trout (*Oncorhynchus mykiss*) from freshwater-to-seawater (SW) for 24h versus 168h as determined via RNA-Seq. Data are presented as changes at 168h relative to 24h. Differentially expressed genes (DEGs) with a Log_2_ fold change ≥ |0.5| between groups and were significantly different following Benjamini-Hochberg false-discovery rate (BH-FDR) correction (p < 0.01) are darkened on the volcano plot and a subset of genes are labelled (see Supplemental File 1 for more details). Note that for visualization purposes the x- and y- axes of the volcano plot are plotted on a Log_2_ and Log_10_ scale, respectively. Significant GO terms were clustered (larger circles) based on overlap between shared genes when analyzing all significant genes, only significant upregulated genes, or only significant downregulated genes. Each smaller circle within the larger circles represents an individual GO term (see Supplemental File 2 for more details). The extent of overlapping genes shared between GO terms is indicated by the intensity of the connecting lines (darker = more shared genes) and significance of each GO term is indicated by the colour of the circle (darker = more significant).

Of the combined 496 distinct DEGs identified between FW- and SW-transferred fish at either 24 or 168 h post-transfer (Fig. 5), most DEGs (96%) were uniquely changed at either 24 h (61%; 161 upregulated and 142 downregulated) or 168 h (35%; 77 upregulated and 95 downregulated). Only 18 DEGs (3.5%) were similarly affected at both timepoints (17 upregulated and 1 downregulated) and 3 DEGs (0.5%) showed opposing patterns between timepoints (1 DEG upregulated at 24 h versus downregulated at 168 h and 2 DEGs downregulated at 24 h versus upregulated at 168 h).

**Figure 5.**
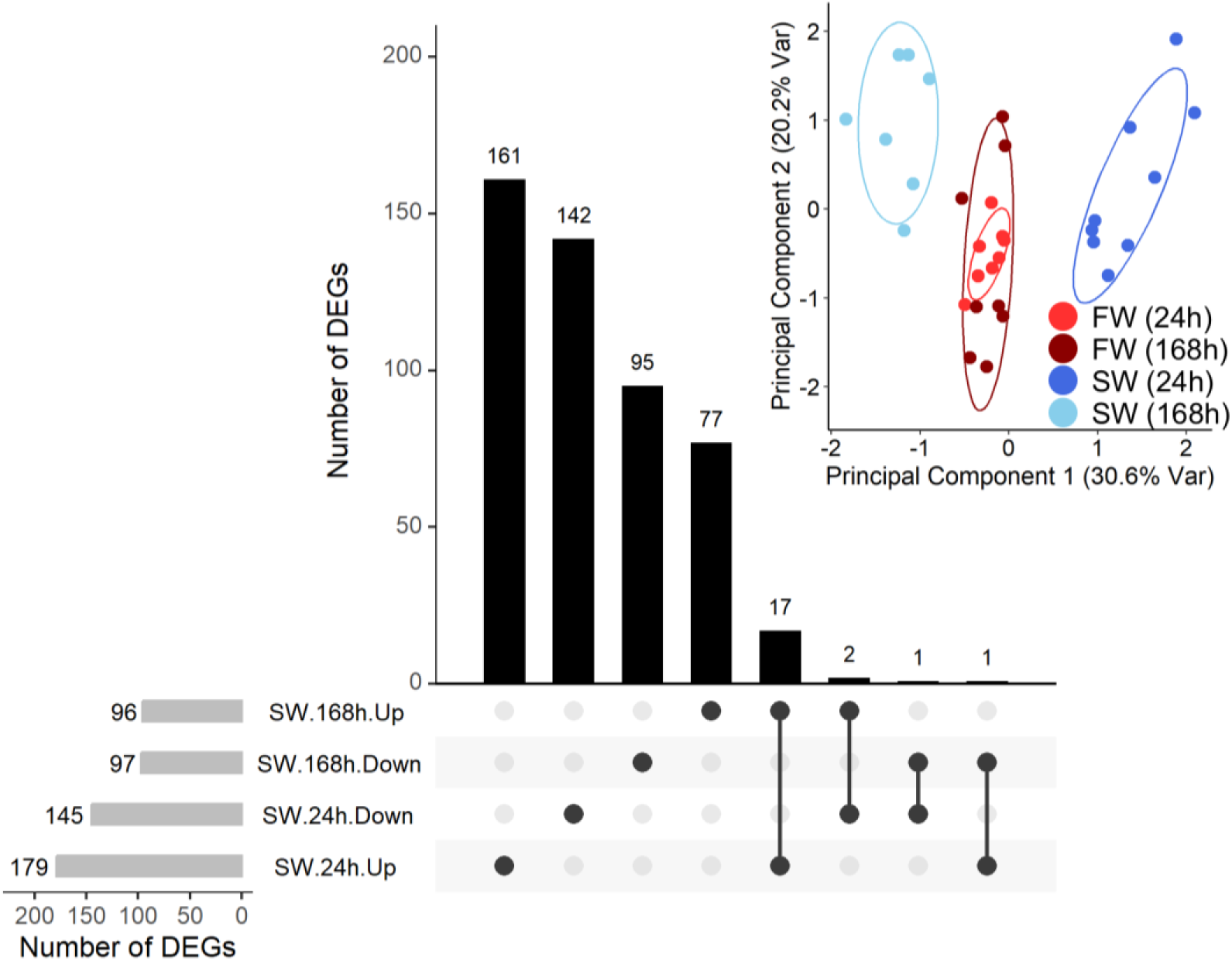
Upset plot depicting overlap between genes that were significantly different in rainbow trout (*Oncorhynchus mykiss*) that were transferred from freshwater-to-seawater (SW) compared to fish that were transferred from freshwater-to-freshwater (FW) for either 24 or 168 h. A biplot depicting overlap between principal components 1 and 2 based on a principal component analysis that included all significant differentially expressed genes (DEGs) is included as an inset.

#### 3.1.3 Presence of endocrine system components in the transcriptome of the CNSS

In addition to detecting components of several peptide (e.g., AVP, CRF, enkephalin, pituitary adenylate-cyclase-activating polypeptide, somatostatin, stanniocalcin, cholecystokinin, neuropeptide Y, and galanin) and steroid (e.g., androgens, corticosteroids, and estrogens) hormone systems that have previously been identified in the CNSS of other teleosts, we also observed components of endocrine systems that have not previously been identified in the CNSS (Table 2). Specifically, we identified components of hormone systems that are broadly involved with calcium maintenance (calcitonin), cardiovascular regulation (natriuretic peptide and tachykinin), growth/osmoregulation [insulin-like growth factor (IGF) and growth hormone], and metabolism (leptin, nesfatin, and thyroid hormone).

**Table 2.**
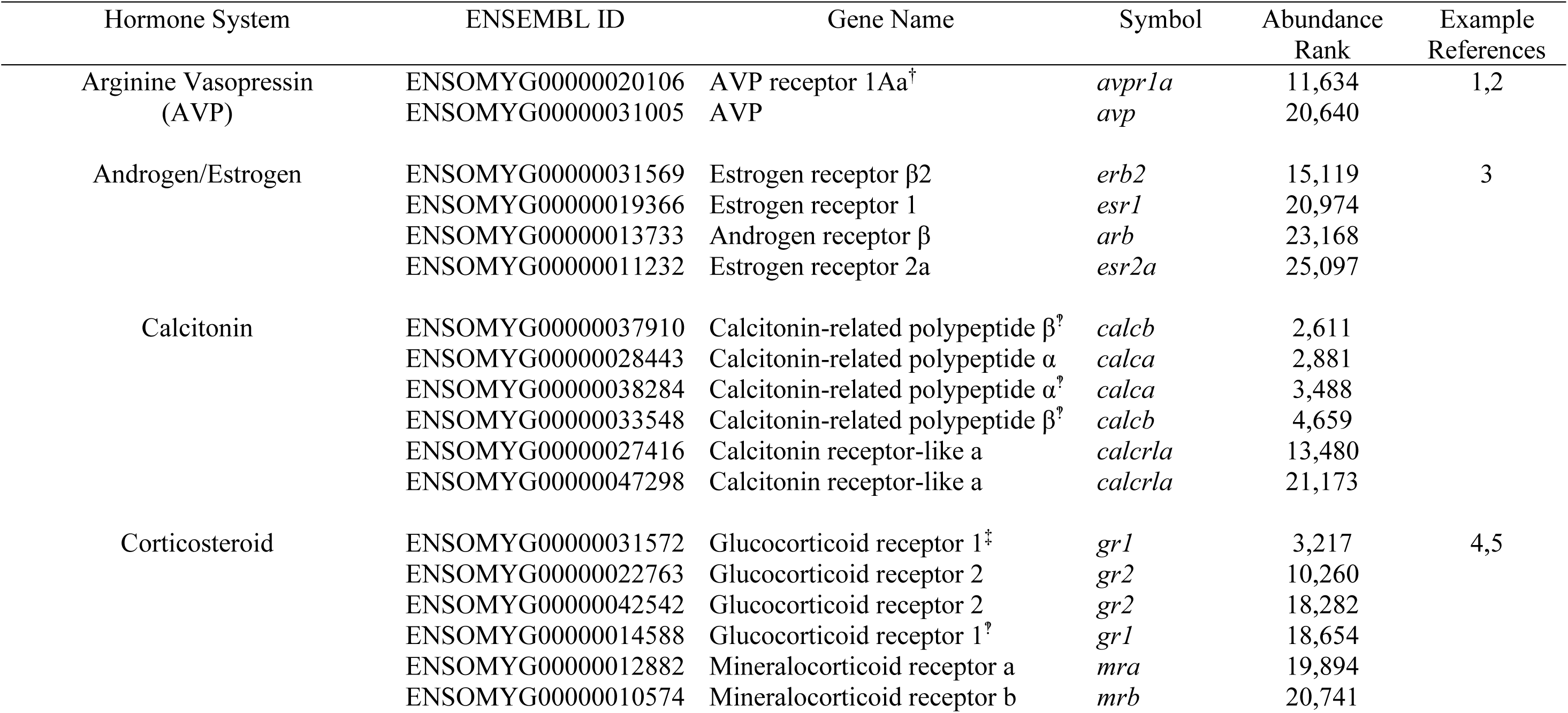
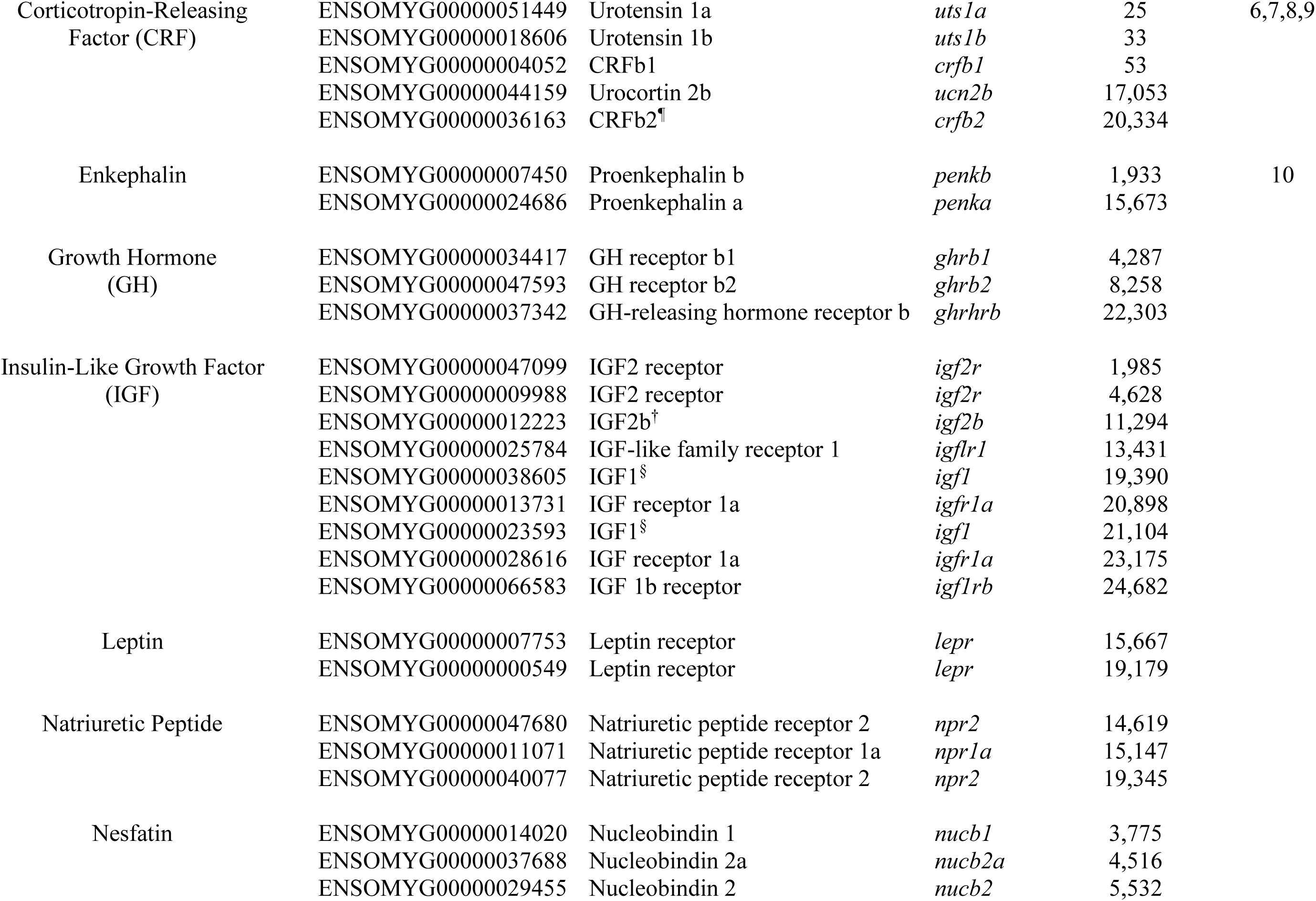
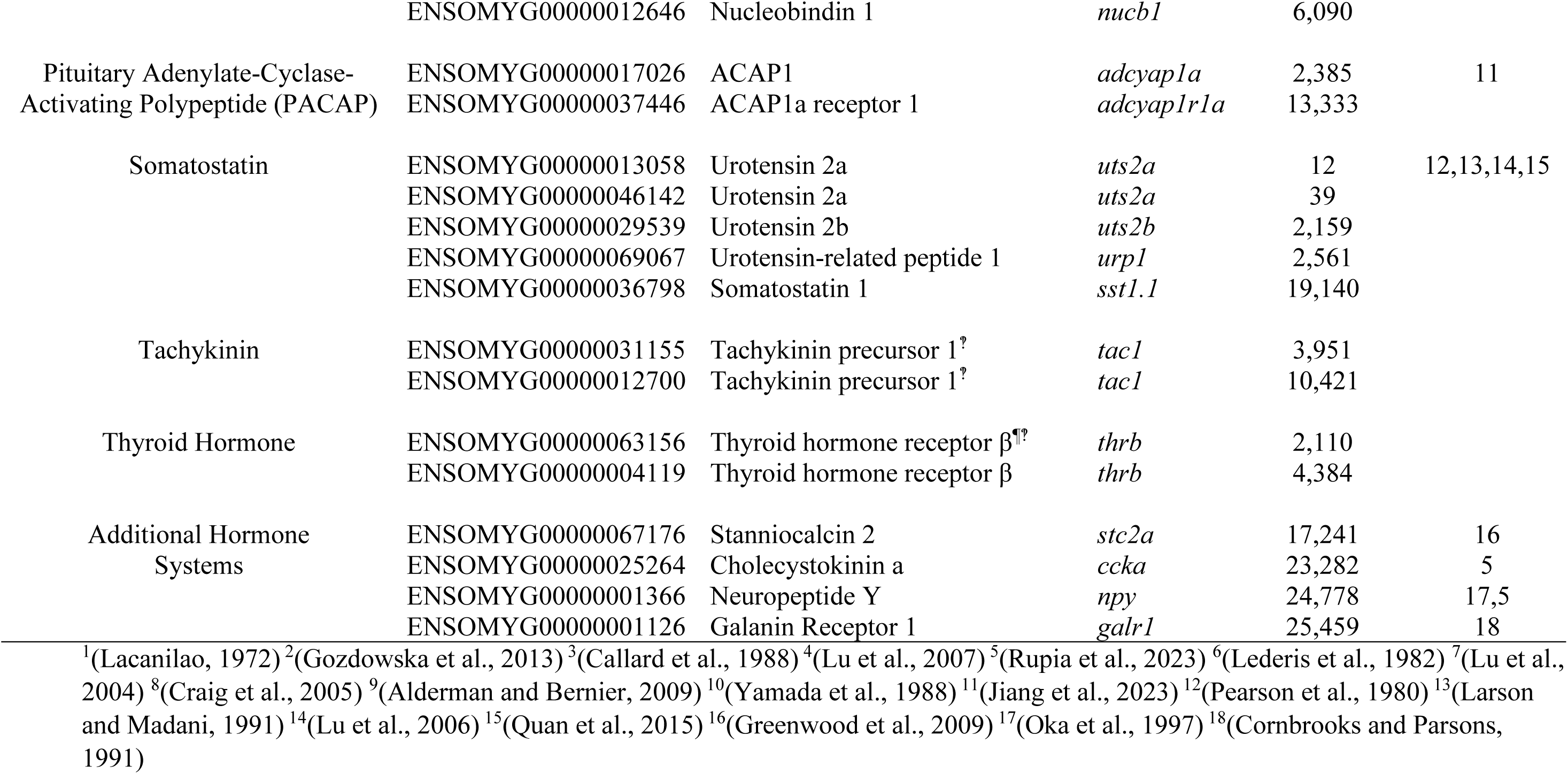
Endocrine-related genes (ligands and receptors) detected in the CNSS of rainbow trout (*Oncorhynchus mykiss*). The relative abundance of each gene is reported as the abundance ranking out of all 25,647 genes detected in the CNSS of rainbow trout in FW (threshold of ≥3 counts per million total reads). Example references are provided in instances where components of each hormone system have previously been identified in the CNSS of a teleost. Genes names are as listed on ENSEMBL except for components of the corticotropin-releasing factor, corticosteroid, and growth hormone systems which were manually annotated in agreement with previous studies of salmonid-specific paralogs for these systems (Mennigen et al., 2022; Romero et al., 2020; Culbert et al., In press). Significant differences (p_FDR_ < 0.05) are indicated using the following symbols: ^†^ = Higher in SW versus FW at 24 h; ^‡^ = Lower in SW versus FW at 24 h; ^§^ = Higher in SW versus FW at 168 h; ^¶^ = Lower in SW versus FW at 168 h; ^‽^ = Higher at 24 h versus 168 h in SW.

Of the endocrine systems present in the CNSS, components of several were affected by SW transfer (Table 2). After 24 h of SW acclimation, levels of AVP receptor 1Aa (*avpr1aa*; +255% compared to FW transferred fish, p_FDR_=0.007), insulin-like growth factor 2b (*igf2b*; +111%, p_FDR_=0.003), and glucocorticoid receptor 1 (*gr1*; -15%, p_FDR_=0.03) were significantly altered. A longer acclimatory period of 168 h in SW caused changes in levels of two paralogs of insulin-like growth factor 1 (*igf1*; +217 and +235%, both p_FDR_=0.04), CRFb2 (*crfb2*; -65%, p_FDR_=0.04), and thyroid hormone receptor beta (*thrb*; -22%, p_FDR_=0.03). Finally, when comparing fish that had acclimated to SW for 168 h versus 24 h, the abundance of two paralogs of calcitonin beta (*calcb*; -28% and -38% compared to SW-transferred fish at 24 h, p_FDR_=0.04 and p_FDR_=0.007), calcitonin alpha (*calca*; -45%, p_FDR_=0.01), glucocorticoid receptor 1 (*gr1*; - 30%, p_FDR_=0.049), two paralogs of tachykinin 1 (*tac1*; -35% and -44%, p_FDR_=0.01 and p_FDR_=0.03), and thyroid hormone receptor beta (*thrb*; -32%, p_FDR_<0.001) were affected.

#### 3.1.4 Effects of FW-to-SW transfer on CRF peptides in the CNSS

In agreement with our RNA-Seq results (Table 2), qPCR assessment of CRF system components in the CNSS of FW-acclimated trout (N=3) indicated that *uts1a*, *uts1b*, and *crfb1* were highly abundant (all amplified at ∼20 cycles; Supp. Fig. 1). Transcripts of *crfbp1* and -*2*, *ucn2a* and -*b*, and *crfr1a* and -*1b* were 8x, 64x, and 1000x less abundant, respectively (with no obvious differences between paralogs), while *crfa1* and -*a2*, *crfb2*, *crfr2a* and *2b*, and *ucn3* were all >4000x less abundant (≥35 cycles). Therefore, we focused all subsequent qPCR analyses on *crfb1*, *uts1a*, and *uts1b*.

Levels of *uts1a* (Fig. 6A; p_group_=0.31, p_time_=0.03, p_group*time_<0.001), *uts1b* (Fig. 6B; p_group_=0.20, p_time_=0.33, p_group*time_=0.002), and *crfb1* (Fig. 6C; p_group_=0.83, p_time_=0.08, p_group*time_=0.003) all showed similar responses following SW transfer. After 24 h, transcript levels were ∼70-170% higher following transfer from FW-to-SW versus FW-to-FW, but levels in FW-to-SW transferred fish declined by 72 h such that they were no longer different than levels in FW-to-FW transferred fish.

**Figure 6.**
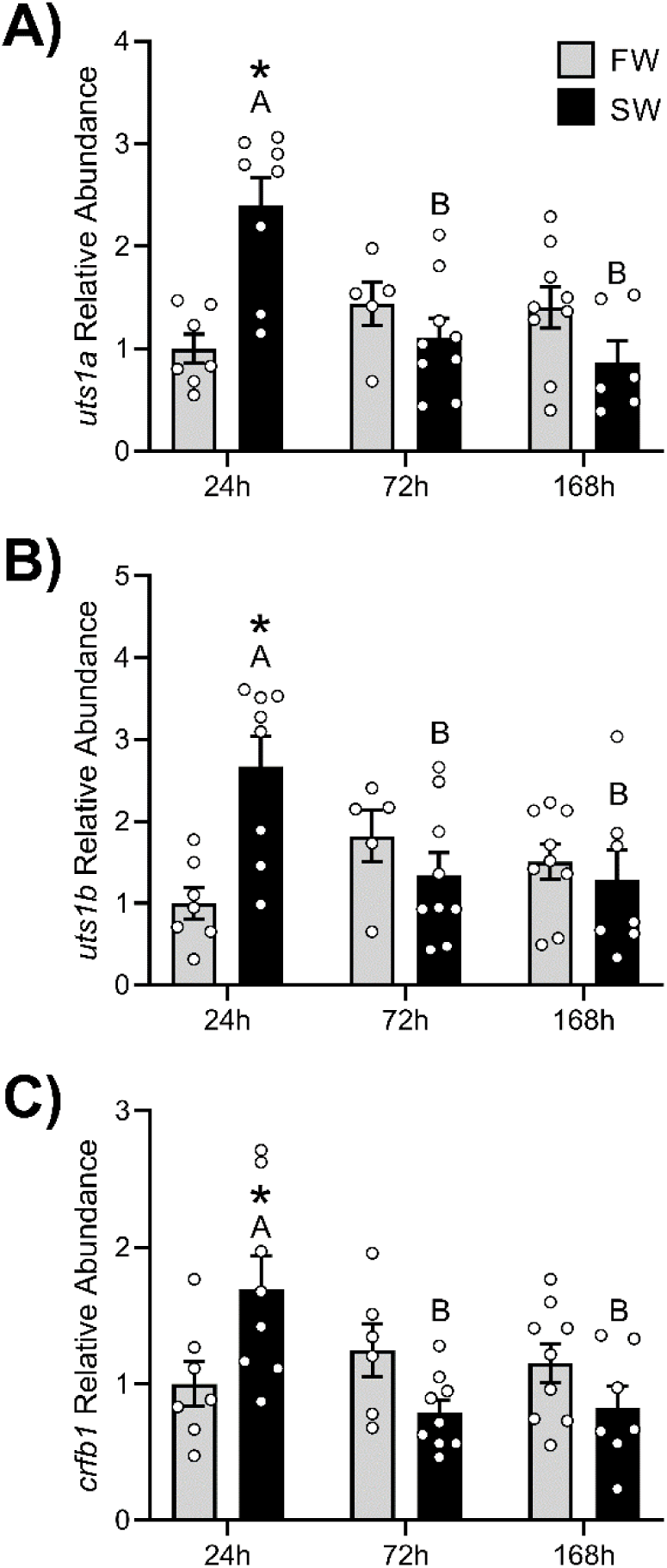
Changes in transcript levels of (A) urotensin 1a (*uts1a*), (B) urotensin 1b (*uts1b*), and (C) corticotropin-releasing factor b1 (*crfb1*) in the caudal neurosecretory system (CNSS) of rainbow trout (*Oncorhynchus mykiss*) that were transferred from freshwater-to-freshwater (FW; grey) or from freshwater-to-seawater (SW; black) for 24, 72, or 168h. Significant differences (p < 0.05) are depicted using either letters (across time) or asterisks (between groups within a timepoint) and data are expressed relative to FW fish at 24h. Values are represented as means ± SEM and individual data points are shown.

### 3.2 Experiment Series 2: Atlantic Salmon

While the following results are focused on transcriptional changes in the CNSS, a full description of changes in plasma osmolality and cortisol levels in these fish has previously been reported (Culbert et al., 2022; Culbert et al., In press). Briefly, plasma cortisol and osmolality values were higher in smolts than parr across all sampling points and parr (but not smolts) had elevated plasma cortisol and osmolality values 24 h after SW transfer. Following transfer from SW-to-FW, plasma osmolality decreased within 24 h while all groups had low cortisol levels (≤ 2 ng mL^-1^).

#### 3.2.1 Seasonal patterns in CRF peptides in the CNSS of parr and smolts

Levels of *uts1a* (Fig. 7A; p_group_<0.001, p_time_<0.001, p_group*time_=0.57), *uts1b* (Fig. 7B; p_group_<0.001, p_time_<0.001, p_group*time_=0.74), and *crfb1* (Fig. 7C; p_group_<0.001, p_time_=0.001, p_group*time_=0.41) were all ∼3‒4-fold higher in smolts than parr through the spring. Additionally, levels increased through the spring in both groups with ∼5-, ∼3-, and ∼2-fold increases in *uts1a*, *uts1b*, and *crfb1*, respectively, when comparing between February and May.

**Figure 7.**
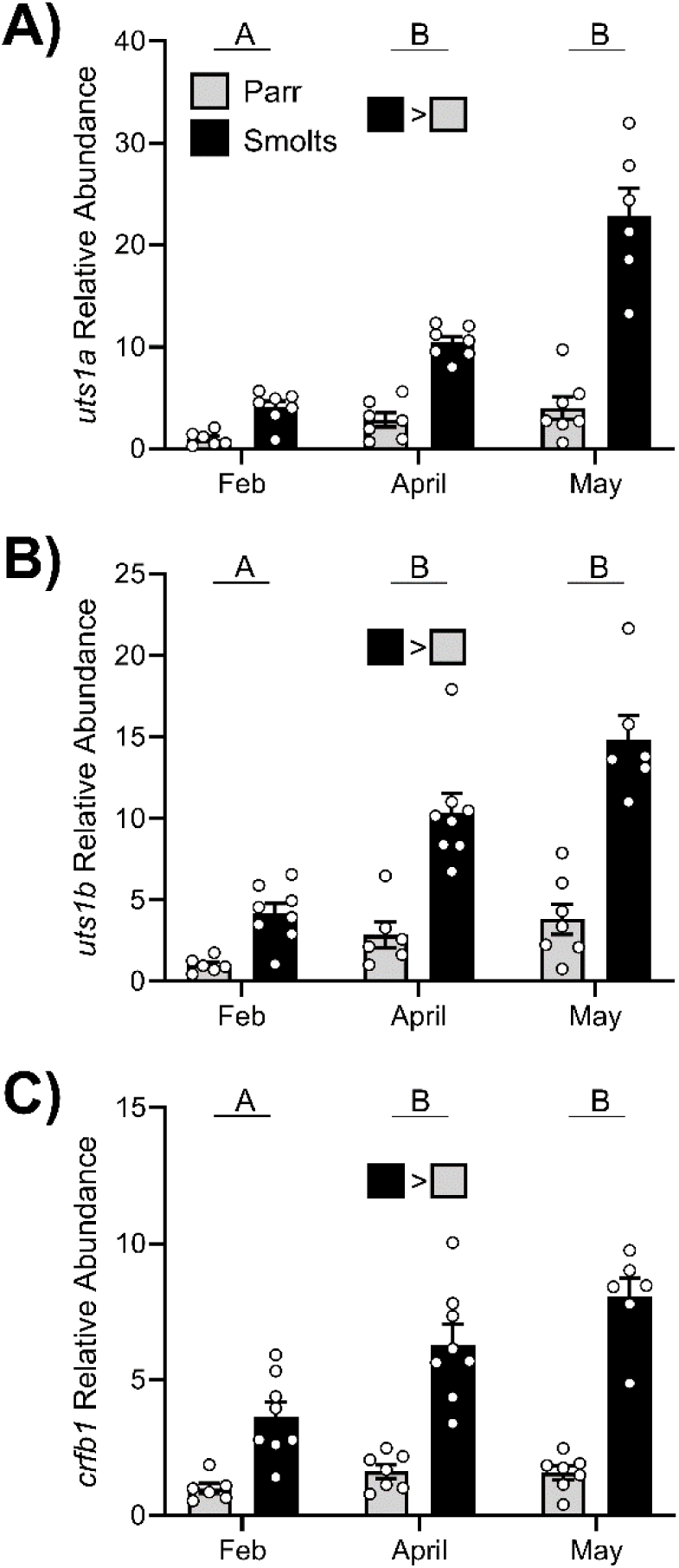
Seasonal changes in transcript levels of (A) urotensin 1a (*uts1a*), (B) urotensin 1b (*uts1b*), and (C) corticotropin-releasing factor b1 (*crfb1*) in the caudal neurosecretory system (CNSS) of parr (grey) or smolt (black) Atlantic salmon (*Salmo salar*) in February, April, and May. Significant differences (p < 0.05) are depicted using either letters (across time; uppercase = within SW, lowercase = within FW, underlined uppercase = overall time effect), filled oversized squares (between groups across all timepoints) or asterisks (between groups within a timepoint) and data are expressed relative to parr in February (Feb). Values are represented as means ± SEM and individual data points are shown.

#### 3.2.2 Effects of FW-to-SW transfer on CRF peptides in the CNSS of parr and smolts

Following transfer from FW-to-SW, levels of *uts1a* (Fig. 8A; p_group_<0.001, p_time_<0.001, p_group*time_=0.15), *uts1b* (Fig. 8B; p_group_<0.001, p_time_<0.001, p_group*time_=0.69), and *crfb1* (Fig. 8C; p_group_<0.001, p_time_<0.001, p_group*time_=0.34) decreased by ∼50‒65% within 96 h in both parr and smolts. However, levels of all peptides remained ∼6‒7-fold higher in smolts versus parr across all timepoints.

**Figure 8.**
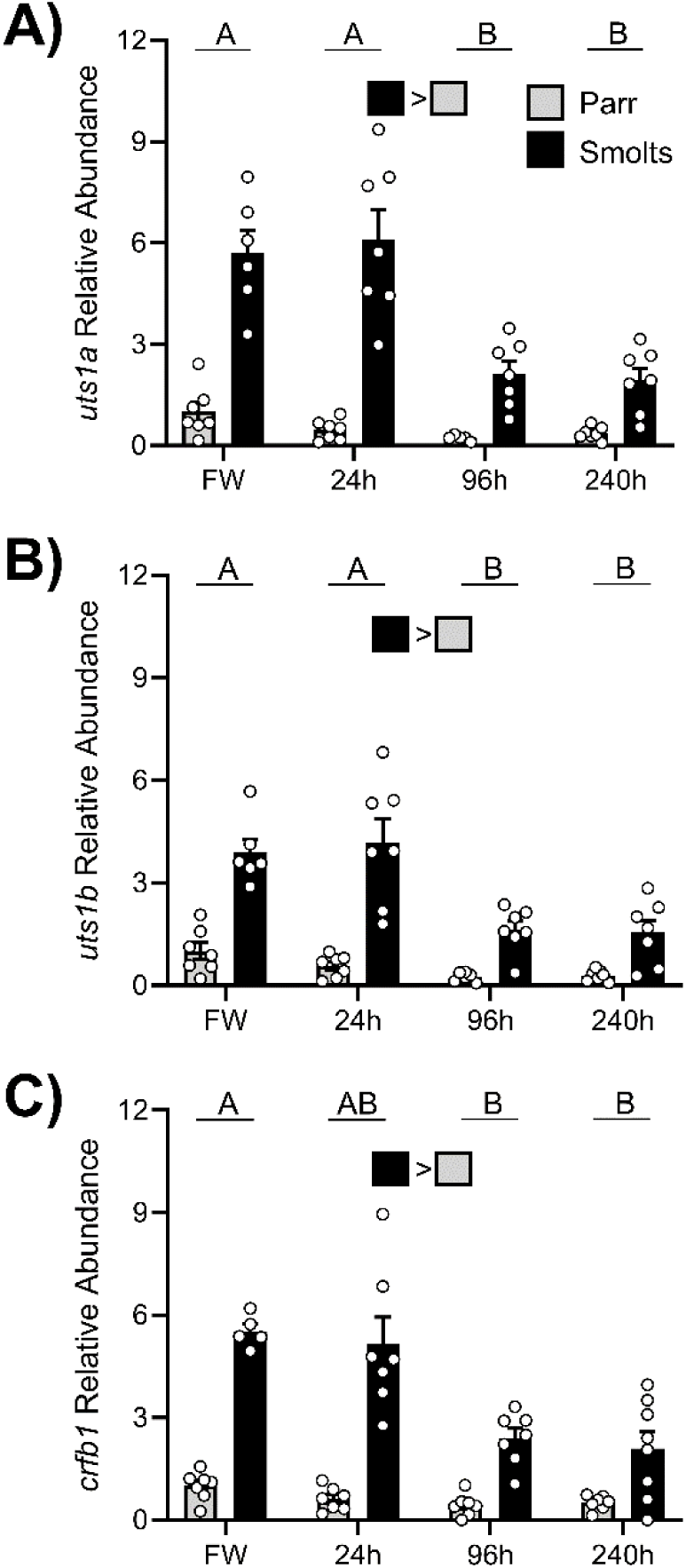
Changes in transcript levels of (A) urotensin 1a (*uts1a*), (B) urotensin 1b (*uts1b*), and (C) corticotropin-releasing factor b1 (*crfb1*) in the caudal neurosecretory system (CNSS) of parr (grey) or smolt (black) Atlantic salmon (*Salmo salar*) that were either sampled in freshwater (FW) 24, 96, or 240h post-transfer to seawater. Significant differences (p < 0.05) are depicted using either letters (across time; underlined uppercase = overall time effect) or filled oversized squares (between groups across all timepoints) and data are expressed relative to FW parr. Values are represented as means ± SEM and individual data points are shown.

#### 3.2.3 Effects of SW-to-FW transfer on CRF peptides in the CNSS of post-smolts

Levels of *uts1a* (Fig. 9A; p=0.17) and *uts1b* (Fig. 9B; p=0.17) did not change following transfer from FW-to-SW, but levels of *crfb1* (Fig 9C; p=0.007) were ∼50% lower 24 h post-transfer compared to fish that were chronically acclimated to either FW or SW.

**Figure 9.**
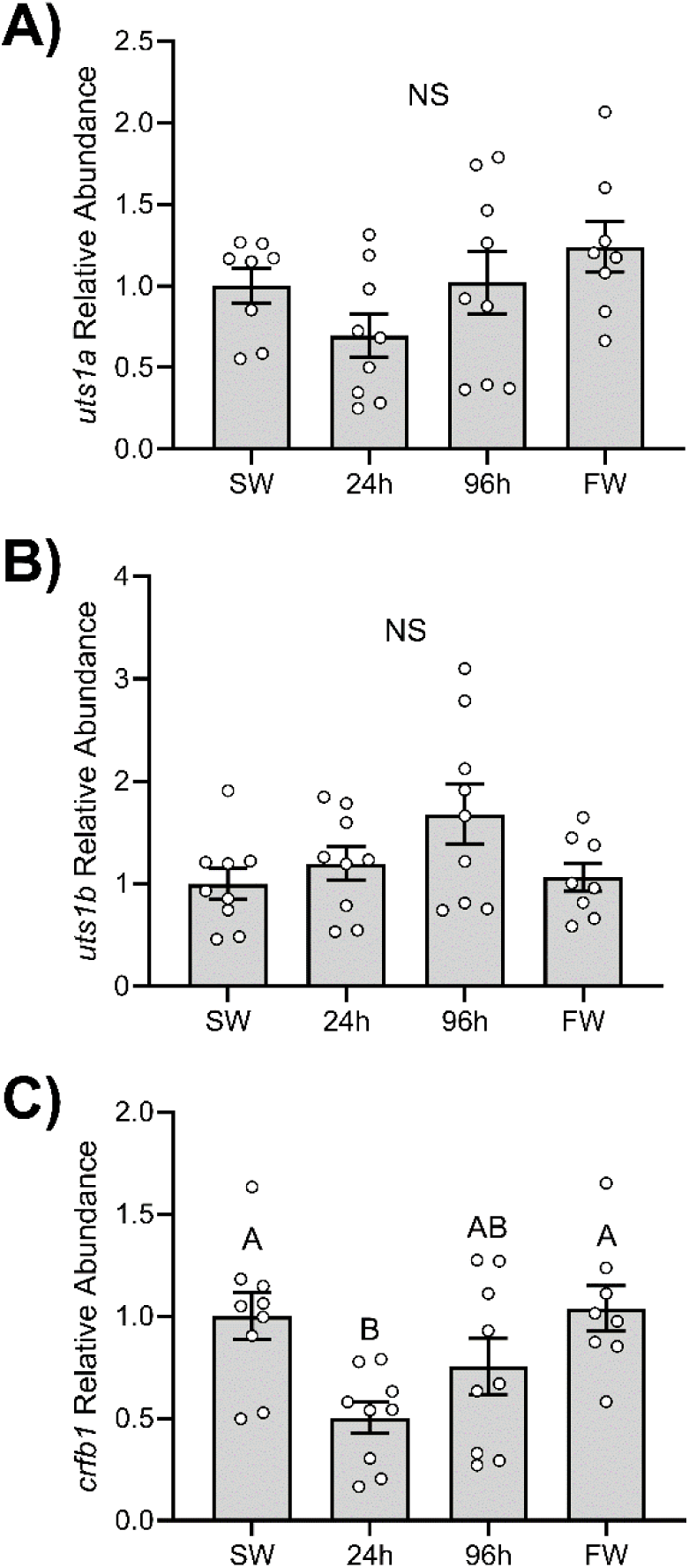
Effects of transferring seawater-acclimated (SW) Atlantic salmon (*Salmo salar*) to freshwater (FW) on transcript levels of (A) urotensin 1a (*uts1a*), (B) urotensin 1b (*uts1b*), and (C) corticotropin-releasing factor b1 (*crfb1*) in the caudal neurosecretory system (CNSS). Significant differences (p < 0.05) are depicted using letters and data are expressed relative to SW-acclimated fish. Values are represented as means ± SEM and individual data points are shown. NS; no significant differences.

## 4. Discussion

While the CNSS has long been hypothesized to serve osmoregulatory roles, our understanding of these roles is limited in part because no study has characterized transcriptome-wide responses by the CNSS following changes in environmental salinity. In the current study, we determined how the transcriptome of the CNSS in rainbow trout—a euryhaline salmonid— changed 24 and 168 h following transfer from FW-to-SW. Plasma osmolality levels were elevated in SW fish at all timepoints—indicating that SW-transferred fish were physiologically perturbed at both timepoints—but plasma cortisol levels had returned to baseline by 168 h, suggesting that fish were beginning to acclimate by this time. Indeed, transcriptional changes in the CNSS at 24 versus 168 h post-transfer support temporal differences in physiological responses following SW transfer. Only 18 DEGs showed conserved responses across both timepoints, and the number of DEGs was far greater when comparing SW-transferred fish at 24 and 168 h (823 DEGs) versus FW- and SW-transferred fish within either timepoint (24 h: 324 DEGs; 168 h: 193 DEGs).

Many of the responses observed 24 h after transfer to SW reflected broad changes in transcriptional processes, including changes in transcription factor signalling (GO cluster: negative transcription regulation), histone regulation (GO cluster: DNA maintenance), and DNA modification (GO cluster: chromosome organization). The only other transcriptomics study which has investigated the CNSS reported similar changes in transcription-related processes in the CNSS of flounder following acute increases in temperature (Yuan et al., 2021), suggesting that this may be a general response of the CNSS following exposure to environmental stressors. Our results are also consistent with transcriptional responses observed in other tissues following SW transfer of salmonids. For example, previous work in Arctic charr (*Salvelinus alpinus*) reported that genes related to transcriptional processes (especially methylation) were upregulated in the gills 240 h after fish had been transferred from FW-to-SW (Norman et al., 2014). Similarly, Harvey et al. (Harvey et al., 2024) found that transcription factor dynamics and chromatin remodeling in the liver of Atlantic salmon changed both during smoltification and following SW transfer. Thus, widespread adjustments in transcriptional processes are clearly an important response while acclimating to SW, as well as responding to stressors more generally.

We also observed several changes in genes related to amino acid transport (GO cluster: membrane transport) and lipid metabolism (GO cluster: lipid biosynthesis) at 24 h, which likely reflects metabolic adjustments associated with SW acclimation. Changes in lipid metabolism— specifically, increased rates of lipolysis and reduced lipogenesis—are one of the primary metabolic changes that occur during smoltification and SW transfer (Gillard et al., 2018; Harvey et al., 2024; Sheridan, 1989), and these periods are also associated with less pronounced changes in amino acid metabolism (Harvey et al., 2024; Sweeting et al., 1985).

Lastly, we detected an upregulation of DEGs related to neurotransmitter transport 24 h after SW transfer, which was primarily related to increased levels of genes that regulate GABA and taurine transport [*slc6a11*, *slc6a6a*, and *slc6a6b*; (Verri et al., 2012)]. While firing rates of Dahlgren cells are regulated by many hormones and neurotransmitters, GABA appears to be one of the major suppressors of Dahlgren cell activity (Lan et al., 2021). Indeed, Yuan et al. (Yuan et al., 2020b) reported that reductions in firing activity of Dahlgren cells in the CNSS of flounder in response to low temperature is primarily mediated by increased GABA signalling. Thus, it is likely that GABA signalling is important for regulating Dahlgren cells activity 24 h post-transfer to SW, but future studies should evaluate changes in other components of GABA signalling pathways (i.e., receptor abundance) to better understand how GABA is affecting the CNSS following SW-transfer. Additionally, increased taurine transport into the CNSS may be an important response following SW transfer since taurine can aid in the maintenance of cellular osmolality and help to minimize oxidative damage (Cheng et al., 2018; Lambert et al., 2015; Schaffer et al., 2000; Thirupathi et al., 2020).

Fewer changes between FW-to-FW and FW-to-SW transferred fish were detected at 168 h post-transfer, with the most notable changes being an upregulation of genes related to protein folding and a downregulation of genes involved in neuron development and differentiation. While upregulation of genes related to protein folding and cellular stress (*calr*, *hsp90b1a*, *dnajb11*, and *hspa5*) likely indicates that SW-transferred trout may still have been acclimating to their new environment even after 168 h, it is less obvious why genes related to neuron development and differentiation might have changed. It is possible that these changes reflect adjustments in the neurosecretory activity of the CNSS (i.e., alterations in Dahlgren cell abundance) since previous work investigating effects of temperature changes on the CNSS of flounder also reported that cellular proliferation was reduced in response to either elevated or reduced temperatures (Yuan et al., 2020a). However, these authors did not observe any changes specifically in populations of Dahlgren cells, suggesting that changes in genes related to neuronal processes detected in the current study are also unlikely to reflect changes in Dahlgren cell abundance. Instead, these changes could reflect reductions in the growth of neurites (progenitors of axons and dendrites) since multiple DEGs in this GO cluster (*slitrk5* and 2 paralogs of *nptna*) are involved in modulating neurite outgrowth and activity (Beesley et al., 2014; Liu et al., 2022); but additional studies are needed to determine what the functional outcome(s) of such changes might be.

Endocrine functions have been associated with the CNSS for 60+ years, yet the suite of hormone systems that are present in the CNSS and/or regulate the CNSS remains unclear. Here, we confirmed the presence of components of many endocrine systems which have previously been identified in the CNSS (e.g., AVP, corticosteroids, CRF, and somatostatin; see Table 2), as well as several hormone systems that have not previously been identified in the CNSS (e.g., calcitonin, growth hormone, IGF, and leptin; see Table 2). Interestingly, despite high abundance of PtHrP components in the CNSS of flounder (Ingleton et al., 2002; Lu et al., 2017), all components of the PTH/PTHrP system that were identified in our RNA-Seq data were very low in abundance (i.e., below the abundance threshold for inclusion in our final gene list). Thus, it is likely that the complement of endocrine systems which are present in the CNSS varies across species and additional work utilizing a diverse range of species is clearly warranted to better establish similarities and differences across teleosts.

In addition to being present in the CNSS, components of several hormone systems in the CNSS were affected by SW transfer. While some of these changes occurred in endocrine system which have previously been identified in the CNSS and have conserved osmoregulatory functions [e.g., AVP, corticosteroids, and CRF; (Cannell et al., 2016; Kulczykowska, 2006; Stengel and Taché, 2009; Takei et al., 2014)], we also detected changes in several hormone systems (e.g., calcitonin, IGF, tachykinin, and thyroid hormone systems) which we do not believe have previously been identified in the CNSS of any fish (Rousseau et al., 2024). Of these novel endocrine systems, changes in calcitonin and IGF systems are particularly interesting since both hormone families have osmoregulatory functions. Calcitonin is involved in calcium regulation (Takei and Hwang, 2016), and calcitonin levels are often acutely elevated following SW transfer (Björnsson et al., 1987; Fouchereau-Peron et al., 1986; Lafont et al., 2006; Najib and Martine, 1996). Indeed, we found that transcript abundance of several calcitonin peptides was higher in SW-transferred fish at 24 h versus 168 h post-transfer. Interestingly, Björnsson et al. (Björnsson et al., 1987) reported that cortisol reduces circulating calcitonin levels in coho salmon (*Oncorhynchus kisutch*). While cortisol levels were 3x higher in SW- versus FW-transferred fish at 24 h, levels of the most abundant paralog of glucocorticoid receptor 1 were 15% lower in SW-transferred fish at this time. Therefore, the observed elevation in calcitonin transcript abundance 24 h post-transfer may be mediated, at least in part, by local reductions in glucocorticoid receptor activity within the CNSS. Like calcitonin, the osmoregulatory actions of IGF-1 (i.e., increased ion secretion and improved SW tolerance) are well established (Mancera and McCormick, 2007; McCormick, 2012); however, far fewer studies have evaluated osmoregulatory functions of IGF-2. In Atlantic salmon and black-chinned tilapia (*Sarotherodon melanotheron*), transfer from FW-to-SW caused transcript levels of both *igf1* and *igf2* to increase in the gills (Breves et al., 2017; Link et al., 2010), but levels of these transcripts were reduced in the brain and liver of tilapia following SW transfer (Link et al., 2022, 2010) suggesting tissue-specific regulation. The current data indicate that transcription of both IGF-1 and IGF-2 is stimulated in the CNSS following SW transfer, but the temporal dynamics of these changes vary. While IGF-2 was upregulated during early acclimation (after 24 h in SW), IGF-1 upregulation didn’t occur until later (after 168 h in SW). In contrast to calcitonin and IGF, there is limited evidence for direct osmoregulatory roles of thyroid hormones in salmonids (McCormick, 2001). Similarly, while tachykinin has many physiological functions (Severini et al., 2002), there is little support for an osmoregulatory role of this hormone family. Therefore, it is unlikely that the observed changes in either thyroid hormone or tachykinin systems are directly associated with osmoregulatory processes. Overall, these data extend our understanding of the importance of endocrine signalling in the CNSS, but additional work is necessary to determine whether these hormones (e.g., IGF and calcitonin) are released into circulation via the urophysis to exert physiological effects on other tissues, or whether the actions of these hormone systems are locally restricted to within the CNSS.

While few significant changes within the CRF system were detected during our RNA-Seq analysis—levels of *crfb2* were 65% lower in SW-to-FW versus FW-to-FW transferred fish at 168 h—we decided to further characterize transcriptional changes in the CRF system of the CNSS using qPCR because CRF peptides were highly abundant in the CNSS and their synthesis has previously been shown to respond to changes in environmental salinity (Craig et al., 2005; Lu et al., 2004). Using this more direct approach, we found that transcript levels of the three major CRF peptide paralogs in the CNSS (*uts1a*, *uts1b*, and *crfb1*) were all 70–170% higher in SW-to-FW versus FW-to-FW transferred rainbow trout at 24 h post-transfer (similar, non-significant responses were also observed in the RNA-Seq analysis). These findings agree with previous studies which have generally shown that transfer of FW fish to SW or marine fish to FW causes acute (≤ 24h post-transfer) changes in protein/mRNA levels of CRF and/or UTS1 (Larson and Madani, 1991; Lu et al., 2019; Minniti et al., 1989), including in rainbow trout (Craig et al., 2005; Larson and Madani, 1996). Interestingly, Craig et al. (Craig et al., 2005) reported that transferring rainbow trout from FW-to-SW caused elevated levels of *crfb* and *uts1* in the CNSS 24, 72, and 168 h post-transfer. It is unclear why our results differ from Craig et al. (Craig et al., 2005), but since body size is positively associated with SW tolerance in many salmonids (McCormick and Naiman, 1984; Parry, 1958), this difference may reflect the size of fish used in our study (∼550g) versus Craig et al. (∼185g). Regardless, because CRF peptides generally promote ion excretion across osmoregulatory epithelia in fish (Chan, 1975; Mainoya and Bern, 1982; Marshall and Bern, 1981), acute activation of CRF peptide transcription following SW transfer likely helps fish develop the appropriate mechanisms needed to minimize the passive influx of ions and promote salt secretion—which is necessary for survival in hyperosmotic environments (Larsen et al., 2014).

In contrast to our rainbow trout results, transcript abundance of CRF peptides decreased when Atlantic salmon parr or smolts were transferred from FW-to-SW. However, this likely reflects prior anticipatory increases in the abundance of CRF peptides which were observed during the spring in both parr and smolts—although, seasonal upregulations were much stronger in smolts than parr—prior to when SW migrations would normally occur. In advance of SW migrations, smolts undergo a variety of physiological adjustments during the spring, including many changes related to osmoregulation, which prepare them to migrate from FW into SW (McCormick, 2012). Based on the current results, it appears that increased activity of the CRF system in the CNSS may be involved in regulating these preparatory changes. Indeed, previous work reported that the cytophysiological activity of the neurosecretory neurons in the CNSS was higher in smolts than parr in coho salmon (Nishioka et al., 1982). Furthermore, Nishioka et al. (Nishioka et al., 1982) also reported that activity in the CNSS of SW-acclimated coho smolts (6-8 months post-transfer) remained higher than FW-acclimated parr, which is consistent with the sustained elevation of CRF peptide transcript levels in smolts even 240 h after SW transfer compared to FW-acclimated parr. While a central role for CRF peptides in the brain have previously been reported in preparation of seasonal migrations in salmonids (Clements et al., 2002; Culbert et al., 2022; McCormick et al., 2019; Ojima and Iwata, 2010; Westring et al., 2008), we believe that the current data are the first report of seasonal changes in the CNSS CRF system of salmonids. Finally, in agreement with the apparent activation of CRF peptide transcription in the CNSS either in response to (rainbow trout) or in advance of (Atlantic salmon) exposure to increased salinity, we also found that levels of *crfb1* transiently decreased 24 h after transfer of SW-acclimated Atlantic salmon into FW. Thus, our data suggest that CRF peptide transcription acutely increases in response to hyperosmotic environments and acutely decreases in response to hypoosmotic environments in salmonids.

Overall, transcriptional responses of the CNSS following transfer from FW-to-SW are temporally dynamic and are associated with changes in several endocrine systems. Additionally, elevated production of CRF peptides by the CNSS, either in response to or in advance of increased environmental salinity, appears to be a conserved response across salmonid fishes. Collectively, these results provide the most intensive investigation of the osmoregulatory functions of the CNSS at a molecular level conducted to date and offer several promising avenues for future studies to evaluate novel physiological contributions of the CNSS.

## CRediT Authorship Contribution Statement

**Brett M. Culbert**: Conceptualization, Data curation, Formal analysis, Investigation, Methodology, Supervision, Validation, Visualization, Writing – original draft, Writing – review & editing. **Stephen D. McCormick**: Conceptualization, Investigation, Methodology, Resources, Writing – review & editing. **Nicholas J. Bernier**: Conceptualization, Funding acquisition, Investigation, Methodology, Resources, Supervision, Writing – review & editing.

## Data Availability Statement

Data will be made available following publication.

## Acknowledgements

We are very appreciative of funds which were provided by Genome Canada and Génome Québec as part of the Genomic Network for Fish Identification, Stress and Health project (GEN-FISH), as well as the technical support provided by Daniel Heath (University of Windsor), Céline Audet (Université du Québec à Rimouski), and Wilian Correa de Macedo and François Lefebvre at the Canadian Centre for Computation Genomics (C3G). We would also like to thank Marcia Chiasson and the staff at the Ontario Aquaculture Research Centre for providing us with the rainbow trout, Chris Wilson and the staff at the Normandale Fish Culture Station for providing us with the Atlantic salmon used in Experiment 3, and Matt Cornish, Mike Davies, and Carolyn Trombley at the Hagen Aqualab for fish husbandry assistance. Finally, we thank Carol Best, Shalya Larsen, Andre Barany, Diogo Ferreira-Martins, Daniel Hall, Jessica Norstog, Amy Regish, and Ciaran Shaughnessy for their assistance with sampling the fish.

