## Supplemental Table and Figure for "Effects of Environmental Salinity on Global and Endocrine-Specific Transcriptomic Profiles in the Caudal Neurosecretory System of Salmonid Fishes"

**Supplemental Table 1**. Number of aligned reads per sample for RNA-Seq analysis on caudal neurosecretory systems (CNSS) collected from rainbow trout (*Oncorhynchus mykiss*) either 24 or 168 h after being transferred from FW-to-FW (FW) or FW-to-SW (SW).

|  | Fish ID | Number of Reads | |
| --- | --- | --- | --- |
| FW |  | 24 h | 168 h |
|  | 1 | 30,969,187 | 44,652,817 |
|  | 2 | 36,797,775 | 32,292,012 |
|  | 3 | 35,814,157 | 41,052,872 |
|  | 4 | 39,130,709 | 29,111,889 |
|  | 5 | 36,734,377 | 27,742,678 |
|  | 6 | 35,552,035 | 37,279,528 |
|  | 7 | 36,293,943 | 29,659,403 |
|  | 8 | 38,900,773 | 29,398,005 |
|  | Mean | 36,274,120 | 33,898,651 |
| SW |  | 24 h | 168 h |
|  | 1 | 41,631,889 | 37,921,975 |
|  | 2 | 35,552,828 | 34,013,244 |
|  | 3 | 37,312,647 | 32,467,065 |
|  | 4 | 31,017,309 | 30,135,233 |
|  | 5 | 36,483,937 | 33,815,889 |
|  | 6 | 35,092,094 | 35,645,927 |
|  | 7 | 36,079,737 | 30,005,528 |
|  | 8 | 29,906,325 |  |
|  | 9 | 33,572,132 |  |
|  | Mean | 35,183,211 | 33,429,266 |

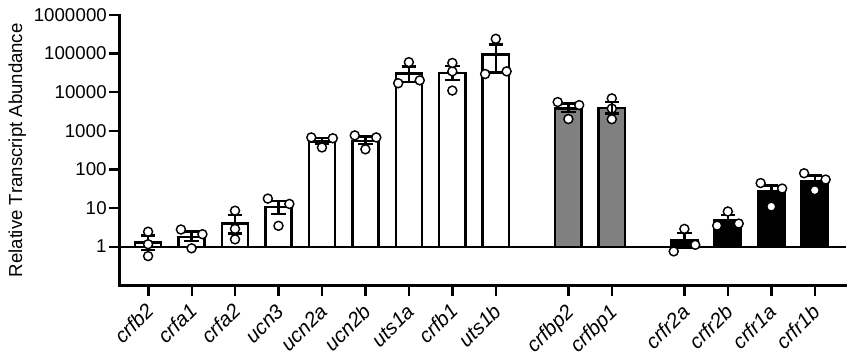
**Supplemental Figure 1**. Relative abundance of individual ligands (white), binding proteins (grey), and receptors (black) of the corticotropin-releasing factor system in caudal neurosecretory system (CNSS) of freshwater-acclimated rainbow trout (*Oncorhynchus mykiss*; N=3). Bars represent mean abundance (± SEM) of each component relative to *crfb2* (the component with the lowest detectable levels). Note that data are plotted on a Log_10_ scale for visualization purposes.
